# A model of egocentric to allocentric understanding in mammalian brains

**DOI:** 10.1101/2020.11.11.378141

**Authors:** Benigno Uria, Borja Ibarz, Andrea Banino, Vinicius Zambaldi, Dharshan Kumaran, Demis Hassabis, Caswell Barry, Charles Blundell

**Author notes:** Equal contribution (Benigno Uria, Borja Ibarz). **Correspondence** Correspondence and requests for materials should be addressed to Benigno Uria and Charles Blundell,).

## Abstract

In the mammalian brain, allocentric representations support efficient self-location and flexible navigation. A number of distinct populations of these spatial responses have been identified but no unified function has been shown to account for their emergence. Here we developed a network, trained with a simple predictive objective, that was capable of mapping egocentric information into an allocentric spatial reference frame. The prediction of visual inputs was sufficient to drive the appearance of spatial representations resembling those observed in rodents: head direction, boundary vector, and place cells, along with the recently discovered egocentric boundary cells, suggesting predictive coding as a principle for their emergence in animals. Strikingly, the network learned a solution for head direction tracking and stabilisation convergent with known biological connectivity. Moreover, like mammalian representations, responses were robust to environmental manipulations, including exposure to novel settings. In contrast to existing reinforcement learning approaches, agents equipped with this network were able to flexibly reuse learnt behaviours —adapting rapidly to unfamiliar environments. Thus, our results indicate that these representations, derived from a simple egocentric predictive framework, form an efficient basis-set for cognitive mapping.

## Introduction

Animals navigate easily and efficiently through the world, learning quickly about new places they encounter^1^. Yet artificial agents struggle with even simple spatial tasks requiring self-localisation and goal directed behaviours. In mammals, the neural circuits supporting spatial memory have been studied extensively. Empirical work has generated a detailed knowledge about populations of spatially modulated neurons – place^2^, grid^3^, and head direction^4^ cells represent body position and direction of facing, while border responsive neurons encode immediate environmental topography^5, 6^. Current thinking sees these networks as components of a cognitive map thought to provide an efficient spatial basis, allowing the structure of novel environments to be learnt rapidly and subsequently supporting flexible navigation, such as short-cuts, detours, and novel routes to remembered goals^2, 7^.

Surprisingly, there are significant gaps in our understanding. In particular, it is unclear how these populations interact to support spatial behaviour and how they are themselves derived and updated by incoming sensory information. Fundamentally, we lack a unifying computational account for the emergence of allocentric (world centred) representations from egocentric (self centred) sensory experience^8, 9^. This poses a problem for neuroscientists, who seek simplifying normative accounts of the brain, and for AI practitioners who aim to build systems with navigational abilities that match those of animals.

Historically, theoretical approaches to this problem largely presented hand-coded designs focused on a single layer of the biological network^10, 11^. A smaller number of models have presented a more extensive description of hippocampal networks^12^, including the process by which representations might be learnt^13^. Notably a potential architecture for ego–allocentric transformation^14^ successfully anticipated the existence of egocentric border responsive neurons^15–17^. Still, in almost all cases existing models are the product of extensive human oversight.

Recently, deep neural networks have provided a normative explanation for the relationship between two allocentric representations – grid patterns emerge when predicting idealised place cell inputs^7, 18–20^– adding to experimental^21^ and developmental^22^ evidence that points to place-cells as an input to downstream grid-cells. Similar approaches have seen considerable success as models of the initial feed-forward processing conducted by the mammalian visual system^23, 24^. However, although impressive, these models have tended to be limited in scope – often being focused on a single biological circuit, taking either idealised neural activity as an input or being constrained to representations bound to a single reference frame.

Here we show that a deep learning model trained to predict visual experience from self-motion is sufficient to explain the hierarchy of egocentric to allocentric representations found in mammals, bridging the gap from visual experience to place cells. In particular, we identify head-direction cells, border cells with both ego and allocentric responses, and place cells. Critically, unlike other recent models of spatial representations^25, 26^, we do so without the need for any allocentric inputs, using only information that is plausibly available to the brain. Thus, our network links this diverse array of cells to a simple predictive framework, it poses hypotheses about the circuit dynamics that produce these responses, and embodies a model linking sensory information, neural activity, and spatial behaviour. Like their biological counterparts, the firing fields of these units are shaped by the surrounding visual cues and boundaries, but retain their characteristics across different environments. When embodied in artificial agents, this redeployment of existing responses supports rapid adaptation to novel environments, enabling effective navigation to remembered goals with limited experience of the test enclosure – demonstrating the flexible transfer of skills at which animals excel but artificial systems have struggled.

### Spatial Memory Pipeline

The hippocampal formation has been characterised as a predictor-comparator, comparing incoming perceptual stimuli with predictions derived from memory^27–29^. With this in mind we trained a deep neural network to predict the forthcoming visual experience of an agent exploring virtual environments. Specifically, we developed a model composed of a modular recurrent neural network (RNN) with an external memory – the ‘Spatial Memory Pipeline’ (see supplementary text, Supplementary Fig 1). At each time-step visual inputs entering the pipeline were compressed using a convolutional neural network and compared to previous embeddings stored in the slots of a memory store, the best matching being most strongly reactivated. The remainder of the pipeline was trained to predict which visual memory slot would be reactivated next (Fig 1A). Predictions were generated solely on the basis of self-motion information, provided differentially to RNN modules as angular and linear velocity, in addition to visual corrections from previous steps. There was no direct pathway from detailed visual features to latter parts of the pipeline – visual information being communicated only by the activation of slots in the memory stores (Fig 1B). Thus the Spatial Memory Pipeline forms a predictive code^30, 31^, anticipating which future visual slot will be activated, based solely on self-motion.

**Figure 1:**
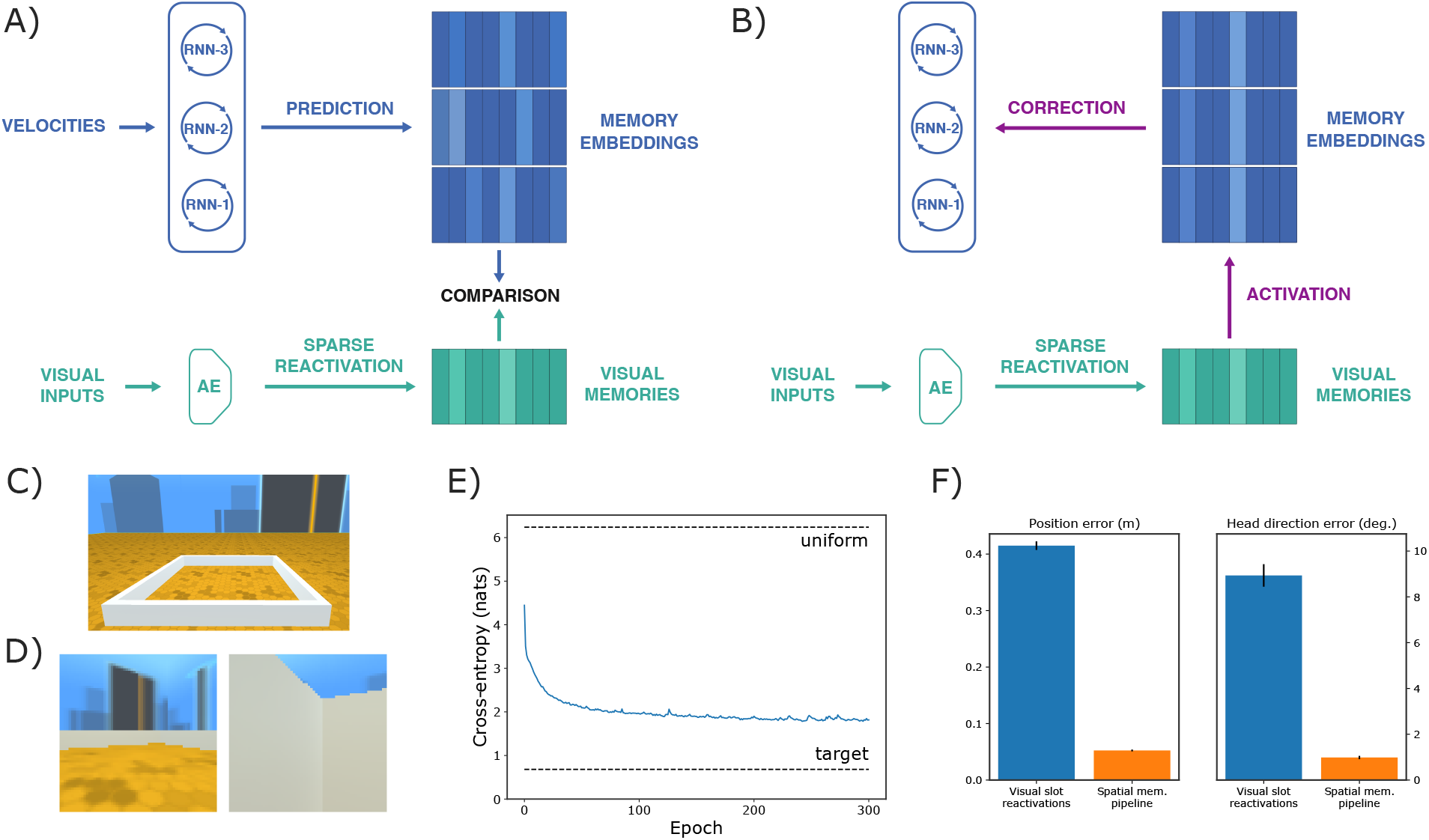
**A, B)** Computational diagrams for the Spatial Memory Pipeline. A) In the prediction step, a set of RNNs anticipate the reactivation of past visual memories by integrating egocentric velocities – AE denotes autoencoder. B) In the correction step, the reactivated visual memories correct the RNNs’ state to reduce accumulated errors. **C)** External view of white square enclosure with distal cues. **D)** Agent’s view from the middle of the enclosure (left) and close to a wall (right). **E)** Prediction loss along training. The model reduces the uncertainty in visual reactivations by 80% with respect to a uniform distribution. **F)** Average decoding error of position and head direction from visual slot activations and spatial memory pipeline RNNs, black bars show one standard error of the mean.

### Emergence of allocentric representations

In our first unsupervised-learning experiment, the Spatial Memory Pipeline was trained in a simulated square environment (2.2 by 2.2 meters) resembling those used in rodent studies – plain white walls with visible distal cues (Fig 1C, D), using a rat-like motion model^32^. After training, the network was able to accurately predict the reactivation of visual memories (80% loss reduction with respect to uniform distribution, Fig 1E), effectively integrating self-motion information to anticipate the visual scene. To understand how the network performed this task we inspected the activity profile of units in the modular RNNs. These units exhibited a range of spatially stable allocentric responses strongly resembling those found in the mammalian hippocampal spatial memory system, including head-direction (HD) cells, boundary vector cells (BVCs), and place cells, as well as egocentric boundary vector cells (egoBVCs) (Fig 2, see Methods). Indeed, in the trained network, the agent’s position and direction of facing could be accurately decoded from the RNN activations (Fig 1F, see Supplementary Methods). These representations were complementary to one another – each RNN module developed distinct response patterns, dependent on the form of self-motion input it received (Fig 2A–D). Similar results were observed in environments with different geometries (see supplementary text, Supplementary Fig 2). The results persisted for different random initializations of network parameters (see Supplementary Results). In contrast, a single RNN receiving all the velocity inputs did not develop the whole range of representations (see Supplementary Results).

**Figure 2:**
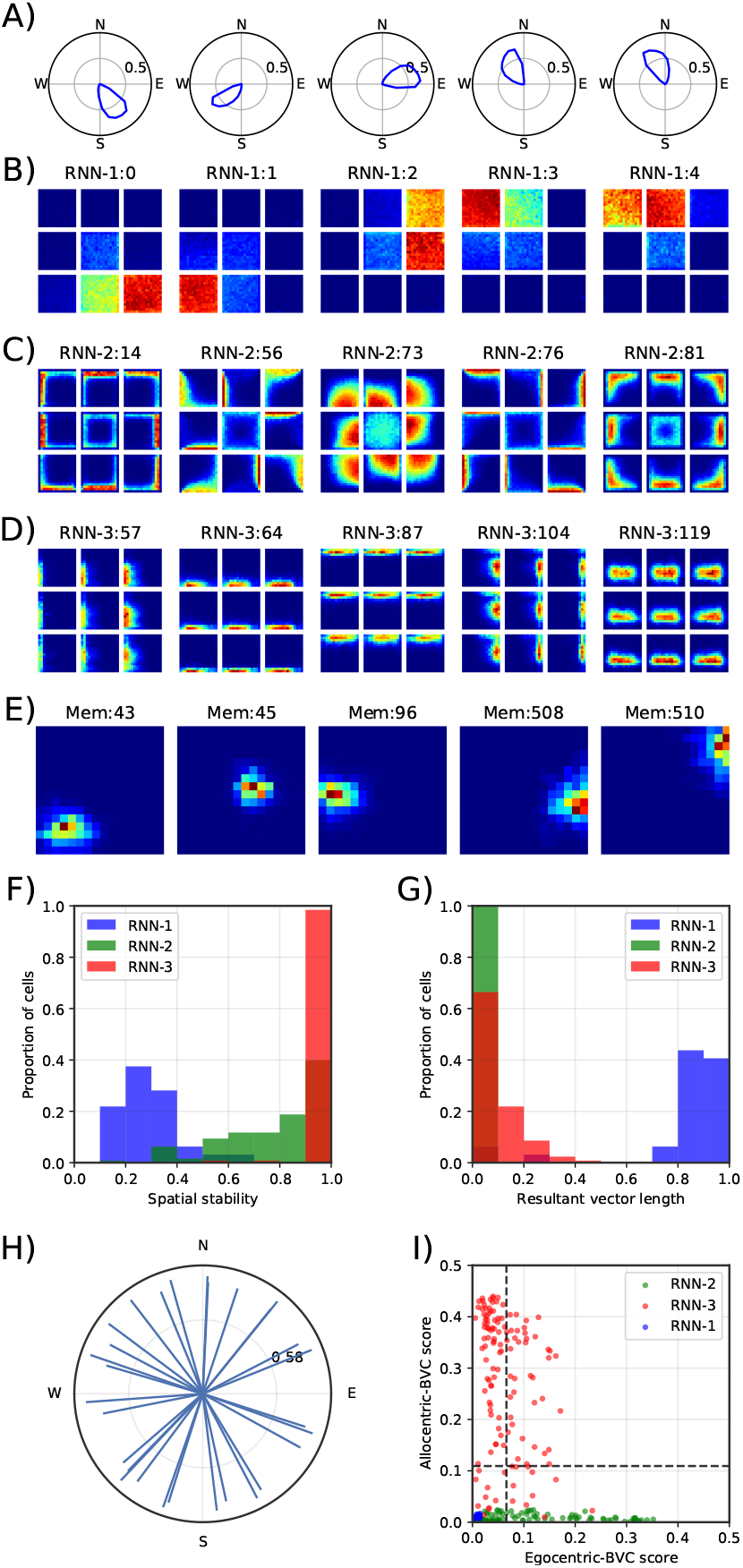
Representations in a spatial memory pipeline trained in the square enclosure shown in Fig 1C. **A)** Polar plot of average activity by heading direction of five units classified as HD cells. **B)** Spatial ratemaps of the units shown in A. Central plot shows average activation for each location across all head directions. The eight plots around each central plot show location-specific activity restricted to times when the heading direction of the agent was in the the corresponding 45° range (e.g. plot located above the central plot shows average activity when the agent was facing in the north direction). **C)** Spatial ratemaps of five units classified as egoBVCs. **D)** Spatial ratemaps of five units classified as BVCs. **E)** Spatial ratemaps of five memory slots reactivated by RNN-3 resembling hippocampal place-cell activity. **F)** Spatial stability of units in each RNN. **G)** Resultant vector length of units in each RNN. **H)** Resultant vector of each unit in RNN-1. **I)** Comparison of egocentric versus allocentric scores for units in each RNN. Dashed lines indicate cell-type classification thresholds.

The majority of units in the first RNN module (RNN-1), which received only angular velocities and corrections from the visual memories, exhibited activity that was modulated by the agent’s direction of facing (Fig 2A). These responses were strikingly similar to those of head-direction cells^4, 33^ – individual units having a single preferred firing direction invariant across spatial locations (Fig 2B). In total 91% (29/32) of units were classified as HD cells (resultant vector length >0.58, 99th percentile of shuffled data, see Methods) with unimodal responses (average tuning width 76.2°) distributed uniformly in the unit circle (Fig 2H, uniform distribution was selected under Bayes Information Criteria (BIC) over mixtures of Von Mises, see Supplementary Methods).

The second module (RNN-2) received angular velocity and speed. Here units exhibited more complex patterns of activity (mean resultant vector = 0.02, 0/128 units classified as HD cells), encoding distances to boundaries at a particular heading (Fig 2C). Thus, they appeared to be similar to egocentric BVCs^15–17^ which are theorised to play a central role in transforming between ego and allocentric reference frames^14, 34^. To quantify this impression we adopted an egoBVC metric applied previously^17^ (see Methods). In this way 52 of 128 units (41%) were identified as egoBVCs (ego-boundary score >0.07, 99th percentile of bin-shuffled distribution, Fig 2I), with the remaining cells displaying mixed patterns of head-direction and egocentric distance to a wall. Rodent egoBVCs exhibit a uniform distribution of preferred firing directions - albeit with a slight tendency to cluster to the animal’s left and right side - while preferred distances have been reported up to 50cm, with shorter range responses being more numerous^17^. Analysis of the model egoBVCs (Supplementary Fig 3A–C) revealed a similar pattern of distances, but a cluster of cells responded to a wall directly in front - plausibly reflecting the monocular input and narrower field of view available to the model (60° vs >180° in rodents) ^35^.

In contrast to the first two RNNs, the third (RNN-3) received no self-motion inputs, thus being dependent upon temporal coherence^13^, and corrections from mispredictions as its sole input and learning signal. Units in this layer were not modulated by heading direction (mean resultant vector = 0.11, 0/128 units classified as HD cells), but were characterised by spatially stable responses (Fig 2F) often extending parallel to particular walls of the environment (Fig 2D). As such, the activity profile of this module was reminiscent of border cells or boundary vector cells (BVCs), which form an allocentric representation of space defined by environmental boundaries^5, 6, 36, 37^. To assess the units’ activity we applied a similar approach to that used for RNN-2 – calculating the resultant vector of the units’ activity projected into allocentric boundary space (see Methods). Applying this measure to RNN-3 confirmed that 84% (108/128) of units were classified as BVCs (BVC score >0.11, 99th percentile of shuffled distribution) – 37% (48/128) met the criteria for egoBVCs, but 37 of those 48 had a higher BVC score than egoBVC score (Fig 2I). The same approach applied to RNN-2 identified 0 of 128 units as BVCs. Analysis of the preferred firing directions and distances of the model BVCs revealed a uniform distribution of directions (Supplementary Fig 3E).

In the brain, BVCs are hypothesised to be a core contributor to the spatial activity of place cells^36^. Consistent with this, BVC-like responses are found in mEC, afferent to hippocampus, as well as within the hippocampus proper^5, 6, 37^. In our model, the allocentric representations of RNN-3 were stored in the external memory slots as a second set of targets - being reactivated at each time step by comparison to the current state of the RNN-3. The activity of these memory slots strongly resembled place cells with sparse, spatially localised responses (average spatial field size of 0.21 *m*^2^ SD:0.26 to a total environment area of 4.84 *m*^2^) that were stable across trajectories, and independent of head direction (see Fig 2E and Supplementary Fig 4).

### Responses to geometric transformations

Simple manipulations of the test environment made without retraining the model, such as rotating distal cues or stretching the enclosure, produced commensurate changes in the receptive fields of spatially modulated cells - resembling the responses of rodent spatial neurons after similar transformations^38, 39^ (Supplementary Fig 5, Supplementary Fig 6, and Supplemental Results). Similarly, an extra barrier inserted into a familiar environment becomes a additional target for BVC and egoBVC responses (Supplementary Fig 7, and Supplementary Results), causing some place fields to duplicate and others to be suppressed^5, 6, 37^.

### Learned attractor dynamics in the head-direction system

In the mammalian brain, head-direction tuning is widely believed to originate from a neural ring attractor which constrains activity to lie on a 1D manifold - this provides a mechanism to integrate head turns when appropriately connected with neurons responsive to angular velocity^40^. In our model, when RNN-1 was instantiated separately and provided only with angular velocities (see Methods), it displayed a single activity bump that closely tracked the apparent head direction for several hundred steps, effectively integrating angular velocity over periods that greatly exceeded the duration of training trajectories (Fig 3A). Projecting the observed activity vectors onto their first two principal components revealed that they resided close to a 1-D circular manifold (Fig 3B), a characteristic associated with linear continuous attractor systems. Indeed, with zero angular velocity inputs, random initialisations of the network state quickly converged to a set of point attractors (Fig 3C–D, see Methods).

**Figure 3:**
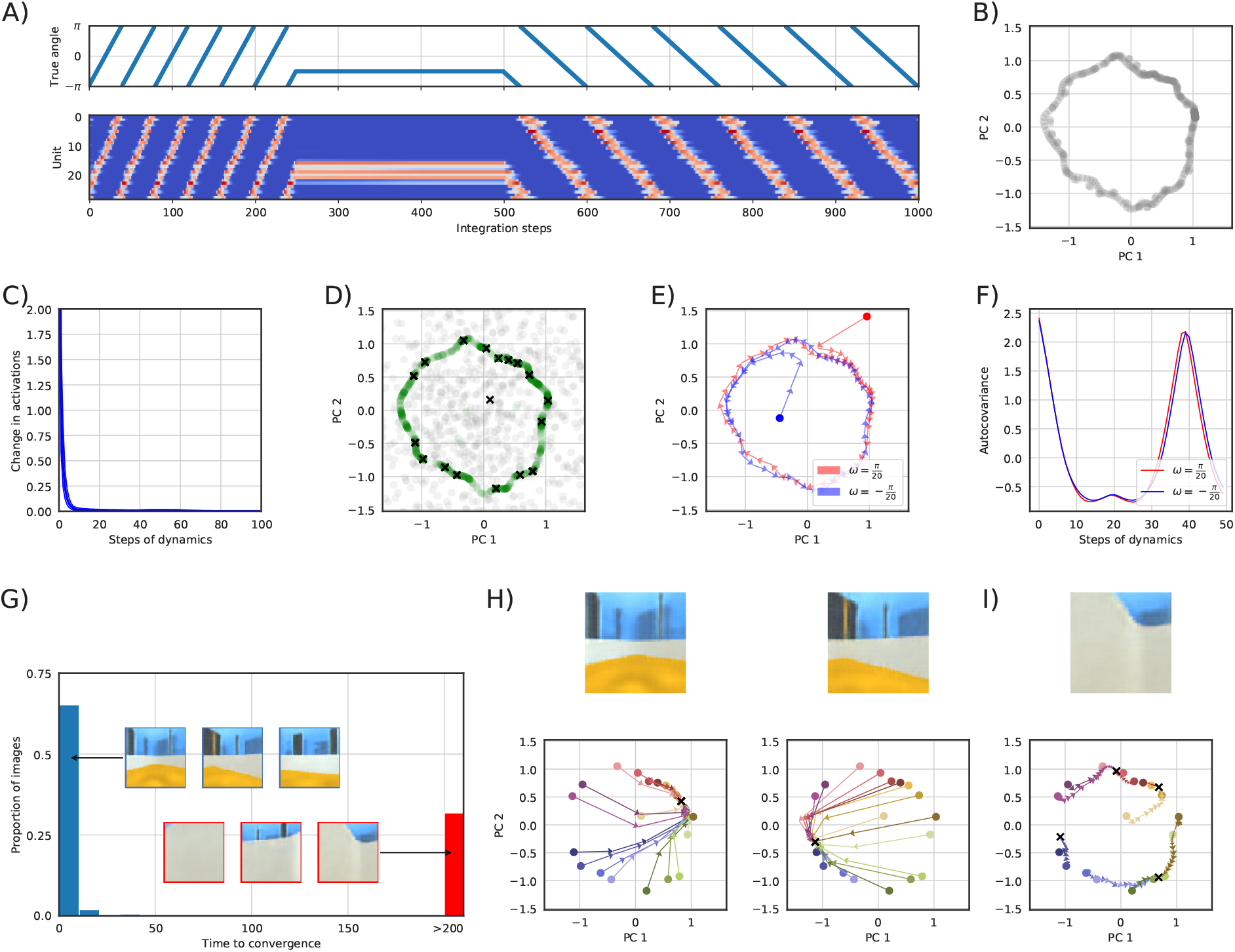
Head-direction cell dynamics. **A)** Activation of each unit over 1000 steps of blind integration. Top: True integrated angle. Bottom: Activation of HD-units in RNN-1 through time (units ordered by the phase of their resultant vectors). For the first 250 steps the angular velocity was set to *π*/20 rad/step, the following 250 steps 0 rad/step, the remaining 500 steps *π*/40 rad/step. **B)** Projection of the 1000 activity vectors shown in A onto their first two principal components. **C)** Euclidean distance between successive states in 1000 trajectories with zero angular velocity and random initialisation (see Methods). Thick blue shows the average over one-thousand different initialisations, the thin lines show the 5-th and 95-th percentiles. **D)** Grey shows the projection of the random initial states used in the simulations of panel C onto the space of panel B. Green shows the state after 10 steps of dynamics with zero angular velocity. Black crosses show the states after 1000 steps. All the trajectories converged to a discrete set of attractor states. **E)** Projection on the space of panel B of two 50-step trajectories with constant angular velocity input, one positive and one negative, each starting from a random initialisation. **F)** Autocovariance of states as a function of time lag computed over 1000 randomly initialised 200-step trajectories with positive and negative velocities. **G)** Histogram of steps to convergence to a single attractor state for 512 correction images. Insets show examples with quick convergence (blue) and some that do not converge to a single state (red). **H)** Several example state trajectories, initialised from the attractor states in panel D (different colours), providing visual inputs with unambiguous cues (top) for 200 steps. Black crosses show the final states. **I)** Same as H with ambiguous visual input of a wall.

In contrast, positive or negative angular velocities drove the state to periodic orbits along 1D cyclic attractors (Fig 3E–F). In the mammalian head direction system these dynamics^4^ are hypothesised to be supported by a double ring network with each ring having counter rotated tuning - a similar solution has been observed in the fly^41^. A strikingly similar connectivity can be learned by simple RNN formulations^42^ (Supplementary Fig 8, Supplementary Results). Angular velocity integration alone is not sufficient to maintain a stable representation of head direction – visual cues are required to reset the activity of units and correct for drift. To investigate how our model incorporates visual information in its representation of heading, we simulated the input of visual corrections (512 images from the training environment) and zero angular velocity (see Methods). The majority of images (352/512) resulted in network dynamics with a single attractor point per image, regardless of the initial network state (Fig 3G). These images typically display unambiguous distal cues (Fig 3H). The remaining images (160/512) did not result in a single attractor point, possibly reflecting the geometrical symmetries of the environment. Interestingly, upon visual examination, these images usually lack distal cues and correspond to views of walls and corners (Fig 3I).

### Spatial characteristics are preserved across environments

Next, we sought to establish how stable the different representations in our model were between environments that changed both in terms of their geometry and visual composition. To this end the network was trained concurrently in three environments inspired by empirical studies – a square, a circle, and a trapezoid, all with distinct distal cues, floor and wall textures (Fig 4A). The same RNNs were used throughout but different external memory stores were provided for each environment. Despite the different training environments, we again found similar proportions of head direction, egoBVC, BVC, and place cells in the network (see Supplementary Results). Indeed, although the environments were visually and geometrically distinct, the classification of units did not materially change between them: 84% (27/32) RNN-1 units were classified as HD cells in all three environments, similarly 78% (100/128) of RNN-2 units where classified as egoBVCs, and 61% (78/128) of RNN-3 units as BVCs. Notably, like biological cells^4^, head direction units maintained their relative angular tuning between environments, while egoBVCs and BVCs maintained their preferred distance to walls (Supplementary Fig 9, see Methods).

**Figure 4:**
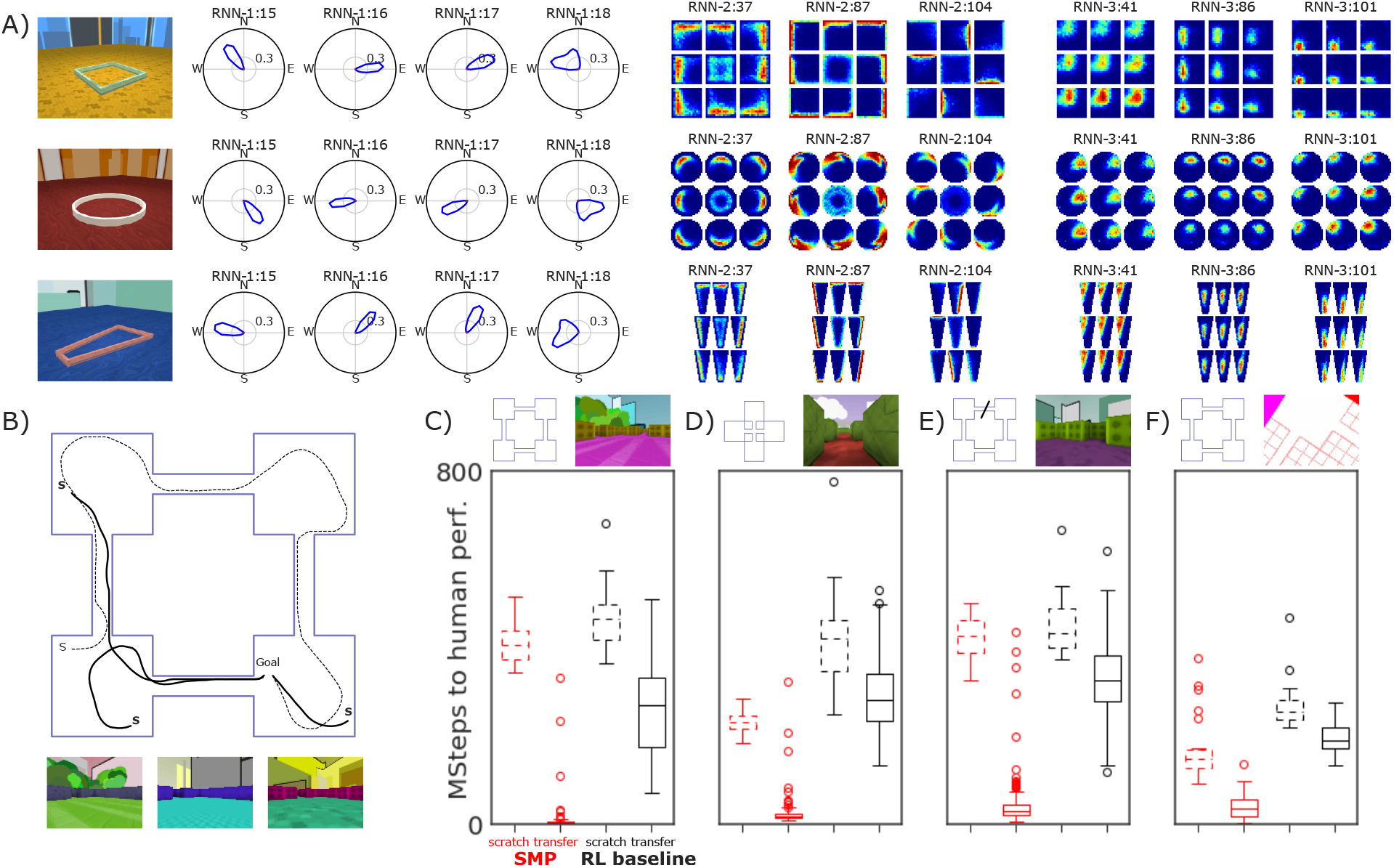
**A)** Stability of spatial representations across three environments, one per row. Left to right: view of the enclosure; allocentric orientation tuning of four units classified as HD cells; spatial ratemap of three units classified as egoBVCs; and spatial ratemap of three units classified as BVCs. Same plotting conventions as Fig 2. **B)** Top, reinforcement learning task in a 4-room enclosure. Spawning points marked with an S. The initial trajectory to the reward is dotted, later trajectories solid. Bottom, visual input from the 3 training enclosures. **C)** Distribution of training times needed to reach human performance in the reinforcement learning task. In red Spatial Memory Pipeline agents, in black baseline agents. Dashed indicates training from scratch, solid indicates transfer to a visually different environment from an agent previously trained in the environments shown in B. On top, schematic of enclosure and example visual input. **D)** Same as C but transfer is to an enclosure with a different layout (cross-shaped). **E)** Same as C but transfer is to a different task, where in each episode a randomly chosen corridor is blocked. **F)** Same as C but transfer is to a conceptual representation of the environment.

### Representation re-use allows transfer of behaviour

In animals, allocentric representations such as head-direction cells and BVCs are rapidly redeployed in novel settings - likely as a result of being constrained to low-dimensional manifolds which decouple the integration of self-motion from high-dimensional perceptual experience^5, 43^. We hypothesised that this self-consistency is an adaptive characteristic allowing spatial behaviour learned in one environment to be quickly transferred to novel environments.

To test this proposal we incorporated the Spatial Memory Pipeline into a deep reinforcement learning agent^44^ (see Methods) trained to find an invisible goal in a 4-room enclosure (Fig 4B), a task inspired by the classic Morris water maze^45^. This task captured two levels of localisation: locally within a room (where in the room?) and globally across rooms (which room?). When the agent reached the goal, it received a reward and was teleported to a random location in the enclosure - to maximise reward it had to reach the goal as many times as possible within each episode. Three visually different enclosures were used simultaneously for training (Fig 4B). As before, separate memory stores were used for each enclosure but RNNs were shared.

Once trained, the agent reached a high degree of proficiency in the task, routinely following the shortest path to the goal (better than human performance). Inspection of the memory pipeline confirmed that the RNNs contained spatial representations consistent across the three visually different environments (Supplementary Fig 12). To test our claim that these allocentric representations provide efficient bases for transfer, the trained agent was placed in a visually different 4-room enclosure (Fig 4C), an enclosure with different room layout and visual aspect (Fig 4D), a variation of the 4-room task where corridors were blocked at random (Fig 4E), and a 4-room enclosure presented conceptually to the agent as a two-dimensional plan view (Fig 4F). In all transfer tasks the agent reached human performance significantly faster than an agent trained from scratch (80x, 12x, 13x and 4x respectively). In contrast, if generic recurrent networks were used instead of the Spatial Memory Pipeline, the agent was still able to learn the initial task but achieved only modest performance gains in transfer (1.7x, 1.5x, 1.3x and 1.3x) (Supplementary Fig 11). Thus, the spatial responses present in the Spatial Memory Pipeline’s RNN support rapid generalisation to novel settings, an ability that is commonly observed in mammals but has been an elusive target for artificial agents.

## Discussion

The Spatial Memory Pipeline described by our model develops an array of sensory-derived allocentric spatial representations strongly resembling those found in the mammalian brain. These representations are learnt solely using a predictive coding principle from egocentric visual perception and self-motion inputs. Like the brain, the network does not have access to allocentric information, and differs markedly from prior work in which brain-inspired agents were limited by the need for allocentric input during training^19, 20, 25, 46^. In contrast, the forms of egocentric representation used here are known to contribute to the activity of spatially modulated neurons in the hippocampal region^47, 48^ while the predictive framework agrees with empirical and theoretical work linking the hippocampus with anticipation of the future^7, 49, 50^. Our core contribution then is the provision of a normative model that explains the appearance of known mammalian spatial representations solely from interaction with the sensory world. This model poses hypotheses about the interactions between these representations, the neural dynamics that generate them, and their role in spatial navigation.

The cell-types that we identified were segregated between different sub-networks, being defined by the forms of information available to them: angular velocity and visual corrections for head-direction cells; angular velocity, speed, and visual corrections for egoBVCs; and only visual corrections for BVCs. External memories reactivated by BVC inputs strongly resembled place cell activity. This framework implicitly captures several properties of the mammalian spatial system - distinct functional populations combining to form a sparse, spatially precise representation of self-location similar to that seen in the hippocampus. Like their biological counterparts, spatial responses in the RNNs were robust to environmental manipulations, responding predictably to the changes made to local cues while retaining their fundamental firing correlates between entirely different enclosures. Our work demonstrates that these representations constitute a flexible and efficient spatial basis, available to be quickly redeployed in a novel setting - sufficient to support rapid transfer learning in a reinforcement learning agent. On the other hand, activity in the memory stores was, by design, entirely distinct between environments, emulating hippocampal remapping, a phenomenon that occurs in response to significant manipulations of environmental cues^51, 52^.

In the case of head-direction cells in the first RNN, the activity of units resulted from learnt attractor dynamics – both in the integration of velocities and in the anchoring to visual cues. Low dimensional manifolds of this form are widely accepted to provide the preeminent account of the biological head-direction system, explaining the fixed relative offset seen in biological head-direction cells as well as in our model^43, 53^. Notably a similar head-direction attractor circuit is instantiated topographically in flies^54, 55^.

A full understanding of the spatial circuits in the mammalian brain will require resolving the functional dependence between different representations. In our model the relationship between the head-direction and BVC systems is determined by their joint association to visual memories (Supplementary Fig 10) - thus they rotate coherently to follow changes in the angular location of a familiar distal cue^56^ (Supplementary Fig 5). Conversely, in different enclosures with non-overlapping sets of visual memories, biological BVCs have been observed to decouple from the head-direction system while retaining their relative tuning^5^. This mechanism of head-direction and BVC dependence contrasts with the hypothesis that postulates BVC activity as the result of direct conjunction of head-direction and egoBVC activations^14, 36^. Hence, new experimental studies on animals jointly recording the head-direction and BVC systems across enclosures that elicit global remapping will be required to ascertain the mechanisms that give rise to BVCs.

In conclusion, we show that an artificial network replicates both the form and function of a biological network central to self-localisation and navigation. A simple training objective is sufficient, in this case, to approximate the development of neural circuits that have been shaped by both selective pressure applied over evolutionary time as well as by direct experience during an animal’s life.

## Methods

### Materials and methods

#### The Spatial Memory Pipeline

The *Spatial Memory Pipeline* consists of two submodules where learning occurs independently. The first level is a visual feature extraction network that reduces the dimensionality of visual inputs. Following visual feature extraction, there is a level of integration through time that encodes estimates of the agent’s location by integrating egocentric velocities. We refer to this integration level as *Memory-Map*. It is composed of a fast-binding slot-based associative memory that plays the role of an idealised hippocampus, and a set of recurrent neural networks that receive as inputs egocentric velocity signals and play the role of an artificial neocortex.

#### Visual-feature extraction

The first level in the *Spatial Memory Pipeline* hierarchy employs a convolutional autoencoder^57^ to reduce the dimensionality of the raw visual input (Supplementary Fig 1A). The autoencoder is composed of an encoder network that transforms the raw visual input, **y**^(*raw*)^ into a low-dimensional visual encoding vector, **y**^(*enc*)^, that summarises the structure of the input. To learn such compressed code, **y**^(*enc*)^ is passed trough a decoder that produces a reconstruction of the original input, 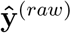. All of the encoder and decoder parameters are optimised to minimise the reconstruction error 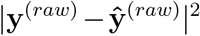. Extended Data Table 4 summarises the autoencoder configuration, see Supplementary Methods for more details.

#### Memory-maps

A Memory-map module consists of two components: a set of *R* recurrent neural networks, {*F*_*r*_}_*r* ∈ 1…*R*_, that integrate egocentric velocities, and a slot-based associative memory, 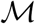 that binds upstream inputs, **y**, to RNN state values:

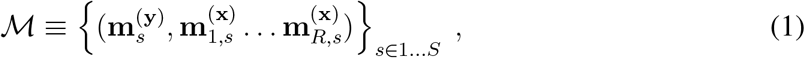

where the variable *s* indexes the different memory slots, while the super-index denotes the type of information (**y** for upstream inputs, and **x** for the state of RNNs).

At each time step, *t*, the current upstream input, **y**_*t*_, reactivates the memory slots with the most similar contents (Supplementary Fig 1B). We formalise this concept using a categorical distribution over memory slots:

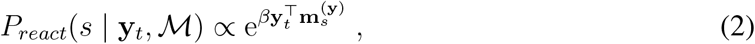

where *β* is a positive scalar that is automatically adjusted to match an entropy target, *H*_*react*_ –enforcing sparse activity (see Supplementary Methods).

At every time step, the states corresponding to each of the predictive RNNs, {**x**_*r*_}_*r*∈1…*R*_ where 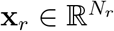, are updated using the current egocentric velocities, **v**_*r,t*_:

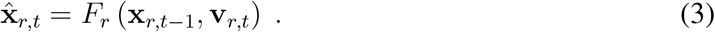

The type of velocity inputs used for each RNN is reported in Extended Data Table 5. For instance, in our unsupervised experiments the first RNN takes as input the sine and cosine of the angular velocity, *ω*, whereas the second RNN takes the same inputs as the first RNN and also the linear speed, *s*. Finally, the third RNN receives no velocity inputs and relies only of temporal correlations.

These RNN states also induce a predictive categorical distribution over memory slots (Supplementary Fig 1B):

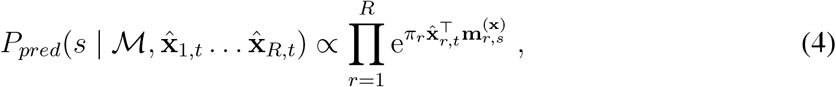

where *π*_*r*_ are positive scalar parameters that determine the relative importance of each RNN in the predictive distribution and its entropy.

Model parameters are optimised to minimise the cross-entropy from this predictive distribution to the distribution of visually driven reactivations:

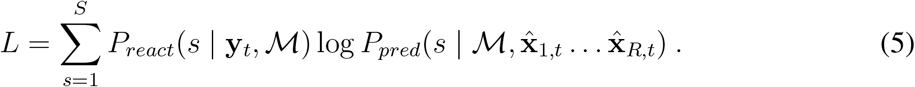

Thus, the model is trained so *P*_*pred*_ anticipates the distribution of memory reactivations *P*_*react*_.

Note that at time *t*, *P*_*pred*_ has not yet received any information from the current upstream visual input, **y**_*t*_, forcing the RNNs to use previous egocentric velocity inputs to produce good predictions.

However, in order for the RNN representations to be allocentrically grounded, and to correct for the accumulation of integration errors, the RNNs must incorporate positional and directional information from upstream visual inputs as well. This correction step should not be performed at every time step, or the integration of velocities would be unnecessary; in our experiments, it was performed at random timesteps with probability *P*_*correction*_ = 0.1. The incorporation of visual information is implemented by calculating a correction code for each RNN:

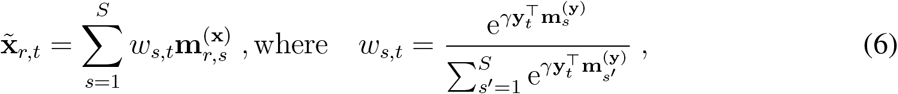

where *γ* is a positive scalar parameter that determines the entropy of the distribution of weights (Supplementary Fig 1C). Each 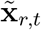 can be thought of as the result of a weighted reactivation of the RNN memory embeddings by the current visual input, **y**_*t*_.

In steps when corrections are provided, these correction codes are input to correction RNN cells, *G*_*r*_, that combine the predictive state with the correction code (Supplementary Fig 1C):

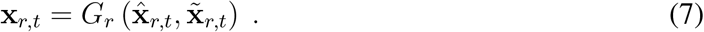

While in steps when no correction is available, the input to the next time step is simply the predicted state (Supplementary Fig 1D):

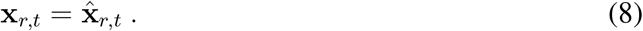

Model parameters 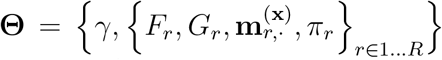 are trained by gradient descent on the prediction loss of equation (5).

Note that the contents of the memory store corresponding to the upstream inputs, **m**^(**y**)^., are not part of the optimised parameters, **Θ**, as they are used to calculate the target distribution. Therefore, in order to fill the slots of 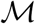 with memories of upstream inputs, at every timestep with small probability, *P*_*storage*_, a slot *s* is chosen at random and assigned 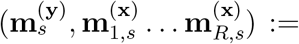 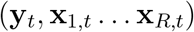.

Due to the sparse activation of memory slots there is low interference between memories, i.e. modifying a 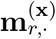. only has a local effect. For this reason these associative memory dings can be optimised at a higher learning rate (see Extended Data Table 2), resulting in fast-binding of new memories once the RNN dynamics have been learned.

#### Architecture used in our experiments

The experimental results in this paper used the architectures described in Extended Data Tables 4-5.

#### Visual autoencoders

The visual-encoding vectors **y**^(*enc*)^ were obtained by inputting RGB images, **y**^(*raw*)^, of the environment to an encoder network with 4 convolutional layers with bias and ReLU output nonlinearities, chained to a flat output layer (Supplementary Fig 1A). The architecture parameters are summarised in Extended Data Table 4. The encoders used for the reinforcement learning experiments had greater capacity than the encoders used for the unsupervised-learning experiments, since they were trained to encode a variety of environments, as explained below.

In our experiments, the visual autoencoders were not affected by the prediction loss in equation (5). Therefore, the autoencoders were trained beforehand, separately from the rest of the model. During the autoencoder training, images were passed through the convolutional layers and flat output layer to produce the vector of visual encodings, **y** ^(*enc*)^. This vector was then passed through a decoder network of de-convolutional layers (with transposed architecture from the encoder but independent parameters) to produce a reconstruction, 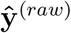, of the input image (Supplementary Fig 1A). All parameters in the autoencoder were optimised to minimise the mean square distance between the input and reconstructed images.

For all experiments, the autoencoders were trained by minibatch gradient descent using an Adam optimiser with learning rate 10^−4^.

For unsupervised experiments, training minibatches consisted of 50 images taken at random from the same trajectories used to train the spatial memory pipeline. Training was complete after 200,000 minibatches. We trained a separate autoencoder for each environment (square, circular, and trapezoid cages). Note that the manipulated environments in Supplementary Fig 5, Supplementary Fig 7, and Supplementary Fig 6, used the same autoencoder as their original, non-manipulated square environment.

For reinforcement learning experiments the training minibatches consisted of 216 images mixed at random from trajectories generated in the training and testing environments using a rat-like motion model (see below). Training was complete after 200,000 minibatches. A separate autoencoder was trained for the 4-room enclosure represented conceptually (Fig 4F), with the same training procedure and architecture.

#### Rat-like motion model

The rat-like motion model used to generate trajectories for unsupervised experiments (and for training the visual autoencoders for reinforcement learning experiments, see above) was based on published work^32^, with parameters specified in Extended Data Table 1. At each time step a linear velocity was generated from a Rayleigh distribution and a rotational velocity from a Gaussian distribution. However, if the trajectory was closer than 3 cm to a wall at an angle narrower than 90°, a deterministic rotation was added that turned it to continue to move parallel to the wall, and the linear velocity was reduced by a factor of 4. Additionally, the trajectory switched between periods of movement (linear velocity Rayleigh parameter 0.13) and periods of pure rotation (linear velocity Rayleigh parameter 0.0). The move and stop periods lasted for an exponentially distributed number of steps with mean 50. The trajectories used to train the spatial memory pipeline were sampled every 7 simulation steps. The angular and linear velocity signals fed to the RNNs in unsupervised experiments were computed dividing the total spatial displacement and angular rotation of the agent over each 7-simulation-step interval by the simulation time of those 7 steps. The corresponding visual embedding came from the image of the environment at the given position and orientation, with the camera 20 cm above and parallel to the ground. Each trajectory consisted of 500 samples, corresponding to 3500 simulation steps.

#### Environment dimensions for unsupervised experiments

For the single-environment unsupervised-learning experiments a square enclosure 2.2 m in side was used. The multi-environment experiments, additionally, included a circular enclosure of radius 1.5 m, and an isosceles trapezoidal enclosure with altitude 4.4 m and parallel sides 2.2 m and 0.75 m long. Single-environment experiments were also carried out in the circular enclosure of 1.5 m radius (see Supplementary Results). The height of the walls in all environments was 0.25 m.

#### Sigmoid-LSTM and Sigmoid-Vanilla

As our aim was to compare the activation of artificial RNNs to firing rate data from biological neurons, we limited the activations of all RNNs to positive values. In order to do this, we modified the commonly used *LSTM* and *Vanilla-RNN* cells by substituting their *tanh* output non-linearity for a *sigmoid*. We called these cells *Sigmoid-LSTM* and *Sigmoid-Vanilla-RNN* respectively. As it learns faster, we used *Sigmoid-LSTM* in all our experiments; with the exception of the supplementary HD double ring analysis (see Methods), where we used a *Sigmoid-Vanilla-RNN* for ease of analysis. In a Vanilla-RNN the mapping from the current state to the next is extremely simple. Namely, the current vector of cell activations, **h**_*t*_, depends on the previous activations, **h**_*t*–1_, and inputs, **x**_*t*_, through equation:

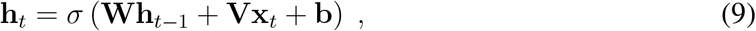

where *σ* is the element-wise sigmoid function, and we refer to **W** as the weight matrix of dynamics and **V** as the weight matrix of inputs.

#### Sparse reactivation

The inverse-temperature parameter in the reactivation of memory slots, *β* in equation (2), is constantly regulated so the entropy of *P*_*react*_ matches the hyperparameter *H*_*react*_ (0.5 nats in unsupervised-learning experiments, 1.0 nats in reinforcement learning). In order to do this, *β* was parameterised as 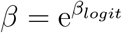 and after every trajectory, *β*_*logit*_ was increased/ decreased by 0.001 when the average entropy of *P*_*react*_ was lower/higher than *H*_*react*_.

#### Training details

The training parameters for the unsupervised experiments are summarised in Extended Data Table 2. The training was implemented with the TensorFlow platform. We used back-propagation through time (BPTT) with an unroll length of 50 trajectory steps; since trajectories consisted of 500 steps, each trajectory produced 10 BPTT unrolls. Batch size was 32, each batch consisting of unrolls from different trajectories.

The single-environment experiments used the Adam optimisation algorithm. Different learning rates were used for the memory slot contents and the rest of the network parameters: since the target distribution over slots was very sparse, we could apply a larger learning rate for the memory slot contents.

For reasons of ease of implementation, the multi-environment experiments used the RMSProp optimisation algorithm. Each of the three environments trained in a separate process, updating the shared parameters synchronously with Distributed TensorFlow. The shared parameters were the weights and biases of the RNNs (both the *F*_*r*_ prediction RNNs and the *G*_*r*_ correction RNNs). The memory contents 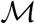 and the entropy-scaling variables *γ* and *π*_*r*_ were separate for each environment.

Similarly to previous work^19^, dropout was used in the RNN outputs when predicting the memory reactivations.

#### Assessment of attractor dynamics in the head-direction system

For the evaluation of integration dynamics in Fig 3A the state of RNN-1 was initialised to 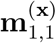, and angular integration (equation 3) iterated for a thousand steps using the angular velocity 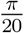 rad/step for 250 steps, 0 rad/step for the following 250 steps, and 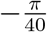 rad/step for the final 500 steps. Visual input was not given to the model. Fig 3A shows the evolution of the RNN state, **x**_1,*t*_, ordering its units by the phase of its resultant vector.

The principal-component space displayed in Fig 3B was calculated by taking the two principal eigenvectors of the covariance matrix of unit activity in Fig 3A. Repeated activity vectors were discarded from the calculation of the mean and covariance matrix.

For the assessment of fixed attractor points in Fig 3C–D, the state of each unit in RNN-1 was independently initialised by drawing a sample from a Gaussian with four times the standard deviation of its activations in Fig 3A. A thousand different random initialisations were used and each run for 1000 steps of angular integration (equation 3) inputting 0 rad/step angular velocity.

For the assessment of periodic orbits in Fig 3E–F, the same one thousand initialisations were used, but each was run for 200 steps of angular integration (equation 3) using either a clockwise 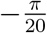 rad/step or counter-clockwise 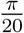 rad/step angular velocity. The 50 first steps of two arbitrary trajectories with opposing velocities were shown in Fig 3E. Each of the two autocovariance curves in Fig 3F was calculated using all 1000 trajectories.

For the assessment of point-attractors under the influence of visual inputs in Fig 3G–I, we took 512 random images from the training dataset, and run the model for 200 steps of prediction and correction. Each of these 200 steps consisted of a step of visual correction, equations 7 and 8, followed by zero angular-velocity integration, equation 3. For every image we run 18 trajectories initialised from each the fixed points found in Fig 3D. We considered that a single fixed-point had been reached at a particular time step if the maximum distance among the 18 trajectories was less than 0.1 in the PCA space. The bottom plots in Fig 3H–I show the trajectory of each of these 18 initialisations for thre examples of visual inputs.

#### Preservation of spatial characteristics across environments

We trained the *Spatial Memory Pipeline* simultaneously in three environments, sharing the RNN parameters across the environments but keeping a separate memory store for each. The separation of memories simulates hippocampal remapping, since there is no correspondence between the slots for the different environments. The RNN units, on the other hand, behaved coherently across the environments, giving further credence to their role as analogues of HD, egoBVC and BVC cells. Supplementary Fig 9 summarises the stability of these representations across the three environments (1, square; 2, circle; 3, trapezoid). The relative preferred angles between HD units in RNN-1 were preserved with great accuracy between environments (Extended Data Fig 9A), i.e., the head direction system rotated as a whole without change in functionality. In contrast, the relative directional tuning of BVC units in RNN-3, although not entirely random (Kuiper test rejected the uniform distribution for the angle differences, *p*-value< 0.01 for all environment pairs), displayed high variability across units (Extended Data Fig 9C). This means that BVC units in our model, under remapping, are not locked to the head direction system, unlike the case without remapping (see rotation manipulations, Supplementary Fig 5). The preferred angles of EgoBVC units in RNN-2 stayed the same in different environments (inter-environment angular differences clustered tightly around zero, Extended Data Fig 9B), as expected for a system of egocentric coordinates. Distance tuning of egoBVCs was very stable across environments (Extended Data Fig 9D), while BVC distance tuning, although significantly preserved (mean across units of the absolute difference between environments significantly smaller than the mean across shuffled pairs, *p*-value< 10^−4^), again showed more variability (Extended Data Fig 9E).

#### Environment for reinforcement learning experiments

We assessed the performance of reinforcement learning agents in the DeepMind Laboratory platform ^58^. The layout of the 4-room enclosures (Fig 4B) consisted of four 5 x 5-tile rooms (1.25 x 1.25 m assuming an agent speed of 15 cm/s) linked by four 5 x 1 corridors. The enclosure was surrounded by a sky-box at infinity - so as to provide directional but not distance information - textured with buildings, clouds and trees, which provided distal cues. There were no special markings on the walls or floors of the enclosure, making all rooms and corridors identical except for their relative orientation with respect to the distal cues. At the start of each episode the agent (described below) was placed in a random location and was required to explore in order to find an unmarked goal, paralleling the task of rodents in the classic Morris water maze. When the goal was reached, the agent received a reward of 10 points and was teleported to a random location in the enclosure. The goal remained fixed for the length of the episode, so a well-trained agent would first explore the enclosure in order to discover the goal location and then, after each teleportation, orient itself to return to the same goal as quickly as possible. The length of the episodes was 3000 agent steps (12000 environment frames, see below).

The agent received observations in the form of camera input (96 x 72 pixels), rotation velocity, and egocentric translation velocity (components parallel and perpendicular to the agent’s orientation).

The agent could start at any position in the enclosure. The action space was discrete (six actions) but afforded fine-grained motor control (that is, the agent could rotate in small increments, accelerate forwards or backwards, or effect rotational acceleration while moving forwards). Agent actions were repeated for 4 consecutive environment frames^44^; one actor step consisted therefore of 4 environment steps.

The visual appearance of the environment was determined by the floor, wall and distal cues. We used 8 different sets of textures, three of them for training and the other five to evaluate transfer. Transfer was evaluated on the same layout as training (Fig 4C), on a cross-shaped layout with 5 x 5-tile rooms linked by two 5 x 1 corridors (Fig 4D), in the same training layout where in each episode one corridor at random was blocked by a wall (Fig 4E), and, finally, on the same training layout provided to the agent not from a first-person camera view but as a top-down schematic representation centered on the agent position and rotated according to the agent’s head direction (Fig 4F). In this schematic representation, each of the 4 boundaries of the layout were filled with a different colour in lieu of distal cues. The image size was 96 x 72 pixels, as in the camera-input experiments.

#### Reinforcement learning agent architecture and training

We used importance-weighted actor-learner agent (IMPALA) ^44^ to learn the task. IMPALA is an actor-critic setup to learn a policy distribution *π* over the available actions and a baseline function *V*^*π*^. Our IMPALA setup consisted of 96 actors, repeatedly generating trajectories of experience, and one learner that used the trajectories sent by actors to learn the policy. At the beginning of each episode actors updated their local policy-network, value-network and *Spatial Memory Pipeline* parameters to the latest learner parameters and, at the end of each episode, sent the trajectory of observations, rewards, states, and policy distributions to the learner. The learner continuously updated its policy, value and *Spatial Memory Pipeline* parameters via back-propagation through time (BPTT) on batches of 64 100-step chunks of trajectories collected from different actors. Both the actor-critic parameters and the *Spatial Memory Pipeline* parameters were optimised simultaneously, but with respect to separate losses. The *Spatial Memory Pipeline* loss is just as described previously in the unsupervised experiments, and determines the gradients for the *Spatial Memory Pipeline* parameters. The actor-critic loss is composed of three terms: a policy gradient loss to maximise expected advantage, a baseline value loss to predict the episode returns, and an entropy bonus to prevent premature convergence. The possible policy lag between the actors and the learner was corrected in the policy gradient and baseline losses with V-trace^44^. We used Adam^59^, a stochastic gradient descent optimiser, for both the *Spatial Memory Pipeline* and the actor-critic loss. A schematic of the agent network is displayed in Supplementary Fig 11A.

The inputs to the actor-critic network were the previous time-step reward and one-hot encoded action, the concatenation of the current outputs of all the *Spatial Memory Pipeline*’s RNNs (current allocentric code), and the the concatenation of the outputs of all the *Spatial Memory Pipeline*’s RNNs observed last time the goal was reached (goal allocentric code, which was set to zero as long as the goal had not been reached in the episode). These inputs were fed to a 256-unit LSTM whose output in turn went through a policy MLP (one 256-unit hidden layer with ReLU activation) to produce the policy, and a value MLP (one 256-unit hidden layer with ReLU activation) to predict the value. Note that the actor-critic network did not receive direct visual input to learn the task - only representations from the *Spatial Memory Pipeline*.

The inputs to the *Spatial Memory Pipeline*, similarly to the unsupervised learning experiments, consisted of egocentric velocities and vision. Because the DeepMind Laboratory agent has inertia, it could shift sideways while moving, even though the action set did not allow strafing. Therefore we provided as inputs not only the translation velocity in the heading direction of the agent, but also the component perpendicular to it. These, together with the sine and cosine of the rotational velocity of the agent, formed the 4-component velocity input vector (as opposed to the 3-component vector used in unsupervised experiments, that lacked a sideways translation component). The visual input was the embedding vector (length 128) of a convolutional autoencoder trained offline to reconstruct images of all the training and transfer environments as explained previously, except the top-down schematic view, for which a separate autoencoder was trained. Compared to the unsupervised experiments, the *Spatial Memory Pipeline* for the reinforcement learning experiments used a bigger RNN-2 size and more memory slots. The hyperparameters of the architecture are described in Extended Data Tables 4-5, while the hyperparameters for training are described in Extended Data Table 3.

In the training phase, each actor was assigned one of the 3 visually different training environments. The *Spatial Memory Pipeline* had correspondingly three sets of memory stores, and experiences coming from each environment were channelled automatically to the corresponding bank. The RNN parameters, in contrast, were shared among all the environments. The agent trained for a combined 2 billion actor steps, then it was tested on the transfer environments. 25 different seeds were used for training, and transfer was evaluated once from each of them.

We evaluated the trained agents in each of the transfer environments separately. In each transfer experiment all the IMPALA actors experienced only the transfer environment, and memory stores were initialised blank. The memory store contents were overwritten at a quicker pace (probability 0.01 per step) at the beginning of the transfer experiments, until full, then continued to be overwritten at the normal rate (probability 0.0001 per step). The rest of the agent parameters started from their values in the trained agent.

#### Reinforcement learning baselines

To support the hypothesis that it is the *Spatial Memory Pipeline* representations that allow the agent to transfer the behaviour to a novel environment with little training, we repeated the transfer experiments replacing the *Spatial Memory Pipeline* with networks of comparable capacity: 1) a generic recurrent network (LSTM) (Supplementary Fig 11B) that integrated visual and velocity inputs, trained with the agent policy loss. 2) A two-layer network (Supplementary Fig 11C) where the first layer consisted of three LSTMs, each integrating the visual and velocity inputs from one of the three training environments, and the second layer consisted of a single LSTM integrating the batched outputs of the first-layer LSTMs; this network paralleled the *Spatial Memory Pipeline* architecture, where each training environment had its own separate memory stores. And 3) a *Spatial Memory Pipeline* identical to the main experiment pipeline, except the visual correction was performed at every time step (*P*_*correction*_ = 1.0); this *Spatial Memory Pipeline* did not need to integrate velocities over multiple agent steps in order to predict the slot activations, and consequently did not develop any useful spatial representations. For replacements 2) and 3), transfer was evaluated only on the same layout - but different textures - as training (Supplementary Fig 11D). All baselines were trained to the same performance as the *Spatial Memory Pipeline* agent before transfer. Replacement 1) had somewhat unstable learning in the top-down schematic representation of the environment (Fig 4F). A smaller learning rate was used in this case, see Extended Data Table 3, since it removed the instability while preserving the average learning curves.

The distributions shown in Fig 4C–F compare the learning time (in millions of actor steps) to reach human performance in each of the 4 transfer settings either training from scratch or transferring from a trained agent, either with the *Spatial Memory Pipeline*-endowed agent or with baseline network 1). Distributions in Fig 4C–E consist of 25 points for the *scratch* box plots (5 environments x 5 seeds) and 125 points for the *transfer* box plots (5 transfer environments x 25 original seeds), distributions in Fig 4F consist of 25 points (1 environments x 25 seeds both for scratch and transfer). For each seed, the time to human performance is measured as the steps required for the average episode reward of the actors to surpass the average human performance and remain above it more than 80% of the following training steps. Human performance was evaluated as the average of 5 episodes played by one of the authors after another 5 episodes of getting acquainted with the task. Computing bootstrapped distributions of the median learning time to reach human performance in each condition yield the following means and standard deviations of the ratios between scratch and transfer medians for the transfer settings in Fig 4C, D, E, F, respectively: for the *Spatial Memory Pipeline*, 85.6 ± 8.9, 11.4 ± 0.6, 13.7 ± 1.0 and 4.4 ± 0.9. For baseline network 1), 1.7 ± 0.1, 1.5 ± 0.1, 1.3 ± 0.1 and 1.4 ± 0.1.

Supplementary Fig 11D–G shows the average learning curves with the standard error of the mean returns across seeds and environments, comparing the *Spatial Memory Pipeline*-endowed agent with baseline network 1). Additionally, Supplementary Fig 11D shows learning curves for baselines 2) and 3) as described above.

The three replacement networks, while able to learn the Morris water-maze task in the training environments, failed to transfer their learned behaviour to visually novel environments (Supplementary Fig 11D). After 50 million steps of transfer training in a novel environment, practically all the *Spatial Memory Pipeline* agents had reached human performance, while none of the agents with generic recurrent networks had, needing a number of training steps for transfer comparable to those needed to learn in the first place (>250 million steps). The *Spatial Memory Pipeline* that was purposefully trained with continuous vision (*P*_*correction*_ = 1.0) to disrupt the development of spatial representations took much longer, demonstrating that it is not an architectural bias, but the spatial representations learned that assist in the transfer of behaviour.

## Data and code availability

The data and code used to generate figures 2 and 3 will be available under the open source Apache 2.0 license at the following URL: https://github.com/deepmind/deepmind-research/tree/master/spatial_memory_pipeline.

There is Supplemental Information that contains additional results, discussion and methods.

## Acknowledgements

We thank Matt Botvinick, Vivek Jayaraman, Tim Behrens, Daan Wierstra, Sergei Lebedev, Christopher Summerfield, Kimberly Stachenfeld, and Charlie Beattie for their valuable advice.

## Author Contributions

Conceived project: B.U, A.B, B.I, C.Ba, C.Bl, D.K, D.H.

Contributed ideas to experiments: B.U, C.Ba, B.I, C.Bl, A.B, V.Z, D.K

Performed experiments and analysis: B.U, B.I, V.Z, A.B

Development of testing platform and environments: B.I, V.Z, B.U, A.B

Wrote paper: C.Ba, C.Bl, B.U, B.I, A.B, D.K, D.H

Managed project:C.Bl, C.Ba, D.H

## Supplemental Information for *A model of egocentric to allocentric understanding in mammalian brains*

This section contains:

- Supplementary Figures
- Supplementary Results

– Emergent representations in a circular environment
– Effects of distal cue rotation on the model’s representations
– Effects of enclosure stretch on the model’s representations
– Effects of barrier insertion on the model’s representations
– Preservation of spatial characteristics across environments
– RNN-1 ring attractor analysis
- Supplementary Methods

– Decoding of position and heading angle
– Quantitative categorisation of spatial representations
– Spatial ratemaps
– Resultant vector of binned data
– Head-direction cells
– Egocentric boundary score
– Allocentric boundary score
– Spatial stability score
– Place cell field characterisation
– Model selection criterion
– Reinforcement learning baselines

## Supplementary Results for *A model of egocentric to allocentric understanding in mammalian brains*

### Emergent representations in a circular environment

The appearance of spatial representations shown in Fig 2 does not depend on the square geometry of the environment. The emergence of such representations is robust to training in enclosures with different environment shapes. To demonstrate this, the model was trained from scratch in a simulated circular environment of radius 1.5 meters, white walls, no internal cues, and distal cues consisting of a city-like scene. The model developed similar representations (Supplementary Fig 2) to those appearing in the square enclosure.

The majority of units in the first RNN module (RNN-1), which received only angular velocities and corrections from the visual memories, exhibited activity modulated by the agent’s direction of facing (Supplementary Fig 2A–B). In total 94% (30/32) of units were classified as head direction cells (resultant vector length >0.48) with unimodal responses distributed uniformly in the unit circle (Supplementary Fig 2H, uniform distribution was selected under BIC over mixtures of Von Mises).

The second module (RNN-2) received angular velocity and speed, together with corrections from the visual memories. Units in this RNN encoded distances relative to boundaries at a particular heading (Supplementary Fig 2C). As in the square arena, they resembled egoBVC responses, with 94% (120/128) of units identified as egoBVCs (ego-boundary score >0.1, 99th percentile of bin-shuffled distribution, Supplementary Fig 2I). None of the 128 units were identified as BVCs. Here cells did not exhibit HD cell-like patterns of activity (mean resultant vector 0.01, 0/128 units classified as HD cells).

The third (RNN-3) received no self-motion inputs, thus was dependent upon corrections from visual reactivations as its sole input and learning signal. Units in this RNN were characterised by spatially stable responses (Supplementary Fig 2F). As in the square enclosure, the activity profile of this module was reminiscent of BVCs (Supplementary Fig 2D). 83% (106/128) of units were classified as BVCs (BVC score>0.11, 99th percentile of shuffled distribution) – 16% (20/128) met the criteria for egoBVCs, but 13 of those 20 had a higher BVC score than egoBVC score. Units in this layer were not modulated by heading direction (mean resultant vector 0.11, 0/128 units classified as HD cells).

### Effects of distal cue rotation on the model’s representations

First, we rotated the visible distal cues by 45° clockwise (Supplementary Fig 5). The preferred orientation of HD units in RNN-1 rotated en masse, tracking the cue rotation the same way that rodent head direction cells are controlled by visual cues (resvec phase rotated by 43° on average, 1° SD) – tuning width was unchanged (average 76.5°, 17.8° SD). The manipulation did not significantly affect egoBVC activity (preferred egocentric directions to wall rotated by 3° on average, 6°SD). The firing correlates of BVCs rotated with the head direction system (35°, 9°SD). The activity field of the place-cell-like memory slots rotated around the centre of the environment (27° on average, 20° SD).

### Effects of enclosure stretch on the model’s representations

We transformed the training environment stretching it by 100% along the horizontal axis to form a 4.4 m by 2.2 m rectangle (Supplementary Fig 6). Again, the response characteristics of HD cells as well as egoBVCs and BVCs were unchanged, and simply extended along the elongated walls like their biological counterparts^5, 6, 16, 17^. Place cell responses closely mirrored those observed in rodents^38^, tending to be stretched along the axis of elongation and in some cases becoming bimodal (Supplementary Fig 6L–O).

### Effects of barrier insertion on the model’s representations

In a similar fashion, we introduced a barrier into the virtual environment (Supplementary Fig 7). The responses of HD cells were largely unaffected, maintaining the same direction of tuning (0° average change in phase, 0.2° SD). Similarly, egoBVCs and BVCs retained their basic firing correlates, responding to the new barrier as they had to the existing walls, resulting in the inception of additional firing fields (Supplementary Fig 7D,H). Biological BVCs and egoBVCs are known to respond similarly; indeed, the predictable response of these cells to extra walls is considered to be a diagnostic feature^5, 6^. Place cell responses were more complex, some – typically those further from the barrier – were unaffected (60%), whereas 22% formed a duplicate field on either side of the barrier (Supplementary Fig 7L–N). Notably, similar outcomes have been reported in empirical studies^60, 37^.

### Preservation of spatial characteristics across environments

To study the properties of the *Spatial Memory Pipeline* representations across environments we trained concurrently in three different enclosures (Fig 4A). In RNN-1, although the preferred firing directions of HD cells changed between environments, the responses of all units were rotated by a similar amount, maintaining the relative angular tuning between cell pairs (resultant vector phase rotation between environments 1 and 2 was 179° on average, 0.5° SD; between environments 1 and 3, 41° on average, 0.6° SD; Fig 4A, Supplementary Fig 9A). The preservation of HD-cell characteristics across environments is consistent with the presence of ring-attractor dynamics, as analysed in single-environment experiments (Fig 3).

The ensemble response properties of egoBVCs in RNN-2 were similarly preserved. Between the three different environments individual egoBVCs maintained their distance and directional tuning, without rotation (mean resultant vector phase rotation 2° on average, 25° SD, between environments 1 and 2; 0° on average, 6° SD between 1 and 3, Supplementary Fig 9B; mean distance tuning difference 0.07 m between 1 and 2, 0.04 m between 1 and 3, *p* < 10^−4^ compared to differences with shuffled units, Supplementary Fig 9D). These representations do not depend on arbitrary distal cues, and are therefore preserved.

The BVCs in RNN-3 tended also to preserve distance tuning (0.19 m mean difference in distance tuning between environments 1 and 2, 0.26 m between 1 and 3, SD 0.22, *p* < 10^−4^ compared to differences with shuffled units, Supplementary Fig 9E), but not directional tuning across environments (phase rotation had SD 69° between environments 1 and 2, and 89° between 1 and 3; Supplementary Fig 9C).

### Robustness to random initialization

The representations exemplified in Fig 2A–E appear independently of the initial values of the parameters of the *Spatial Memory Pipeline*. In all of 20 experiments run with identical hyperparameters but different random parameter initializations, HDs, egoBVCs and BVCs appeared in numbers similar to those reported in the main text. BVCs always appear exclusively in RNN-3. HDs appear typically in RNN-1 as described (>31 out of 32 units classified as HD in 12 out of 20 experiments), but sometimes they are also found in RNN-2. Specifically, in 7/20 experiments HDs appeared exclusively in RNN-1, as in the main text example; in 5/20 they appeared both in RNN-1 and RNN-2; and in the remaining 8/20 they appeared exclusively in RNN-2. In such cases, all RNN-1 units were classified as egoBVCs. Since angular velocity information is available to RNN-2, it is not unexpected that HDs can be learned there too, and, to the effects of prediction optimization, separating RNN-1 and RNN-2 is unnecessary. Doing so, however, allows us to simplify the analysis of the attractor dynamics of the HD system.

Similarly, the experiment described in Fig 4A and Supplementary Fig 9 was typical, but in some of the experiments HDs appeared also in RNN-2. Specifically, across 15 multi-environment experiments (using 5 different random seeds and 3 different RNN learning rates, 10^−3^, 3 . 10^−4^, and 10^−4^), 8 had HD cells purely in RNN-1, as in the experiment shown, while the other 7 had a significant number of them in RNN-2 instead.

### Single-RNN ablation

Splitting the model into three separate RNNs with different velocity inputs is important to obtain all of the representations. If a single RNN with both linear and angular velocity input is used, with the same number of units as the total in our 3-RNN model (288), approximately 1/3 of the units will be HDs and 1/3 will be egoBVCs; the remaining 1/3 do not satisfy any of the classification criteria. Remarkably, no BVCs appear. Conversely, if the single RNN receives no velocity inputs, approximately 60% of the units are classified as BVCs, and nearly 30% of units classify as egoBVCs, but no HDs appear. Thus, having both an RNN with velocity inputs and one without is necessary to produce HDs, egoBVCs and alloBVCs in the model. As noted earlier, the split between RNN-1 and RNN-2 is not strictly necessary but simplifies the analysis of the HD system.

### RNN-1 ring attractor analysis

To understand the mechanism by which our model tracks head direction, we repeated the experiment shown in Fig 3 substituting the efficient but complex LSTM integrators with Vanilla-RNNs. In Vanilla-RNNs each activity state is a simple function of the previous state and inputs, thus is amenable to a mechanistic analysis^42^ (see Supplementary Methods). As expected, this new experiment also developed head-direction cells in RNN-1, all 32 units being classified as head direction cells (resultant vector length >0.4, 99-th percentile of shuffled data, Supplementary Fig 8A). Like the LSTM-based model, the Vanilla-RNN effectively integrated angular velocity over many time steps (Supplementary Fig 8B).

The simplicity of Vanilla-RNNs allowed us to examine the connectivity that supported angular velocity integration, revealing a striking similarity with that found in the fly^41^ and hypothesised for mammals^40^. The weight matrix of dynamics, **W**, when displayed ordered according to each units’ preferred firing direction (Supplementary Fig 8C), resembled a circulant matrix, with a diagonal band of excitatory connections surrounded by diagonal bands of inhibition. That is, the connectivity between units forms a ring with local excitation and long-range inhibition. The angular-velocity tuning of the cells, calculated as the arc-tangent of the 2-dimensional vector of control weights, **V**, that receive as input the cosine and sine of the angular velocity, showed a clear split of the units in two groups, 18/32 units being activated by positive angular velocities (ccw-cells) and 14/32 activated by negative angular velocities (cw-cells) (Supplementary Fig 8D). Plotting the weight matrix of dynamics separately for each group revealed the mechanism by which angular velocity was integrated (Supplementary Fig 8E–F). Each cell preferentially excited other cells offset around the ring in the same direction as their angular-velocity tuning, an asymmetry that became more obvious when the average weights to units with different relative firing directions were calculated (Supplementary Fig 8G).

## Supplementary Methods for *A model of egocentric to allocentric understanding in mammalian brains*

### Decoding of position and heading angle

For decoding of position and head direction (Fig 1D) we trained a single-layer MLP (256 hidden units with tanh non-linearity) to predict either the true (x, y) position of the agent in the environment, or the cosine and sine of its true head direction, from either the vector of log probabilities of activation of the visual slots or the concatenation of all the RNN outputs. We used RMSProp as optimiser (learning rate 10^−4^, decay 0.9, no momentum) with square loss. The decoder was trained on 10 million batches of 25 frames sampled at random out of 300,000 frames from the same trajectories used for unsupervised training of the Spatial Memory Pipeline. The decoding error shown in Fig 1D for position is the mean Euclidean distance over the whole training set between the (x, y) output of the decoder and the true position. For head direction, it is the mean absolute difference between the true head direction and the arc-tangent of the cosine and sine prediction of the decoder. Error bars show the standard error of the mean calculated as the standard deviation of 100 bootstrapped means computed from 5000 decoded samples.

### Quantitative categorisation of spatial representations

Where possible we have used the same functional characterisations of cells used for rodent data. Resultant vector lengths of allocentric direction-of-facing were used to characterise head-direction cells^61^. Resultant vector lengths of egocentric-boundary ratemaps were used to characterise egocentric boundary cells (egoBVCs) ^62^. We used the same procedure with allocentric-boundary ratemaps to detect allocentric boundary cells (BVCs). Our results were based on datasets of 800 trajectories each consisting of 500 timesteps.

Our simulation datasets of 400,000 timesteps correspond to much longer duration than is common in rodent studies. For this reason we avoided using thresholds based on random shifts of the data, as this results in very low thresholds due to a null hypothesis that removes most of the dependency of activations from any environmental features. We used a bin-shuffling procedure that results in much more stringent, and qualitatively pleasing, thresholds. Thresholds were calculated as the 99-th percentile of resultant vector lengths for 1000 bin shuffles for each unit. All units from all RNNs were used to calculate the threshold of each cell category.

### Spatial ratemaps

We measured and plotted the dependence of a cell’s activity on the agent’s location by using spatial ratemaps. We partitioned the environment into a grid of 14cm by 14cms square bins aligned with the cardinal directions. The spatial ratemap for a cell is the matrix of its average activity when the agent is located inside each of these bins.

When we aimed to show the dependence of cell’s activity on both spatial location and head-direction, the per-octant spatial ratemaps were calculated and plotted surrounding the spatial-ratemap. These are eight spatial ratemaps where the data is restricted to times when the agent’s head direction is contained in each of the eight intervals of 45° centered at the cardinal and inter-cardinal directions.

### Resultant vector of binned data

Resultant vectors were calculated as a single complex number from mean activities binned by angle:

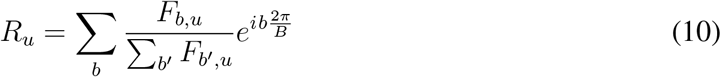

where *F*_*b,u*_ is the average activity of unit *u* for the *b*-th bin out of a total of *B* bins.

The length of the resultant vector corresponds to the modulus of *R*_*u*_ and serves as a statistical measure of circular concentration (inverse variance). While the argument of *R*_*u*_ is a statistical measure of preferred direction of activation.

### Head-direction cells

Head direction cells characteristically fire in a small range of allocentric direction. This preference for a single direction can be measured by the length of the resultant vector of activations.

In order to calculate the resultant vector of each unit, we partitioned allocentric directions into 20 bins. The angular ratemap, *F*_*b,u*_, of the *u*-th unit, with instantaneous activity *a*_*u,t*_ at time *t*, was calculated as:

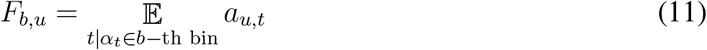

where *α*_*t*_ the agent’s direction of facing at time *t*, and *α*_*t*_ ∈ *b*-th bin if 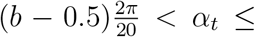 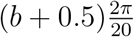.

The tuning-curve width for units classified as head-direction cells was calculated as 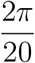, the width of a single bin, times the number of bins where the mean activity was greater than 0.5 of the greatest mean-bin activity for that unit.

### Egocentric boundary score

Egocentric boundary cells were characterised by the resultant vector of a the egocentric-boundary ratemap (EBR) ^62^. The EBR uses the agent’s position and direction of facing as frame of reference and computes the average instantaneous activity when a boundary is present at a particular distance and egocentric direction. In our unsupervised experiments the EBR was calculated for bins of 4° by 2.5cm. The maximum distance considered was half of the maximum distance between opposing walls. Examples of EBRs can be seen in the middle row of Supplementary Fig 3A and D.

The length of resultant vector of the EBR (averaged over distance) was used to detect egocentric boundary cells. For a cell classified as an egoBVC, the angle of the resultant vector was taken as its preferred direction and the distance of the maximum activity along this direction in the EBR as its preferred distance.

### Allocentric boundary score

We characterised allocentric boundary cells using an allocentric-boundary ratemap (ABR). In the ABR the agent’s position is used as frame of reference but not its direction of facing, the relative angle to boundaries is locked to the allocentric north direction. The ABR computes the average instantaneous activity where a boundary is present at a particular distance and allocentric direction. In our unsupervised experiments the ABR was calculated for bins of 4° by 2.5cm. The maximum distance considered was half of the maximum distance between opposing walls. Examples of ABRs can be seen in the bottom row of Supplementary Fig 3A and D

The length of resultant vector of the ABR (averaged over distance) was used to detect allocentric boundary cells (BVC). For a cell classified as a BVC, the angle of the resultant vector was taken as its preferred direction and the distance of the maximum activity along this direction in the ABR as its preferred distance.

### Spatial stability score

Where reported, the spatial stability of a unit was measured as the Pearson correlation of its spatial ratemap for two halves of the data. In our experiments each half of the data was comprised of 200,000 data points, which under the motion model used would amount to 3000 seconds for each half.

### Place cell field characterisation

The activity field of a cell was calculated using the following procedure on its spatial ratemap: (1) a Gaussian filter of radius 1 bin was applied, (2) bins with value higher than half the maximum value were considered part of the activity fields, (3) fields were segmented using the `skimage.measure.label` function, (4) fields smaller than 3 adjacent bins were discarded, and (5) the attributes of each field (position, size, and eccentricity) were calculated using the `skimage.measure.regionprops` function with the original ratemap as `intensityimage` parameter. We used version 0.14.0 of `skimage` throughout.

### Model selection criterion

Where a probability distribution is reported as fitting data (e.g. the distribution of HD angles, or the angular preferences of egoBVCs) we used the Bayesian information criterion to compare several alternative distributions. This criterion penalises the log-likelihood of the model, *L*, by a term that depends on the number of parameters, *k*, and number of data points, *n*:

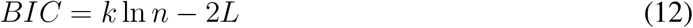

A lower BIC signals a better model fit.

### Reinforcement learning baselines

Supplementary Fig 11A depicts the architecture of our agent as described in Methods. The baselines we used to compare performance in the water maze transfer tasks are the following:

- An agent with the same architecture, receiving visual corrections at every time step (instead of every 10 steps on average). This is the baseline labelled *SMP transfer no integration* in Supplementary Fig 11D.
- An agent where the *Spatial Memory Pipeline* was replaced by a generic recurrent network (LSTM) (Supplementary Fig 11B) integrating visual and velocity inputs. The size of the LSTM output was 928 units, to make it the same as the concatenation of all the RNN outputs in the *Spatial Memory Pipeline*. Unlike the memory pipeline, the LSTM did not have a separate unsupervised loss for training, but was trained directly from the gradients of the policy loss.
- An agent where the *Spatial Memory Pipeline* was replaced by a two-layer network of LSTMs (Supplementary Fig 11C), where the first layer consisted of three LSTMs (output size 512 units), each integrating the visual and velocity inputs from one of the three training environments, and the second layer consisted of a single LSTM (output size 928 units) integrating the batched outputs of the first-layer LSTMs. This network paralleled the *Spatial Memory Pipeline* architecture, with the first-layer LSTMs playing the role of the per-environment memory banks and the second-layer LSTM the role of the memory pipeline RNNs. For transfer, the first layer was replaced by a single 512-unit LSTM, and the rest of parameters were kept from the trained agent. As with the single-layer LSTM baseline, the LSTMs did not have a separate unsupervised loss for training, and were trained directly from the gradients of the policy loss.

---

**Supplementary Figure 1:**
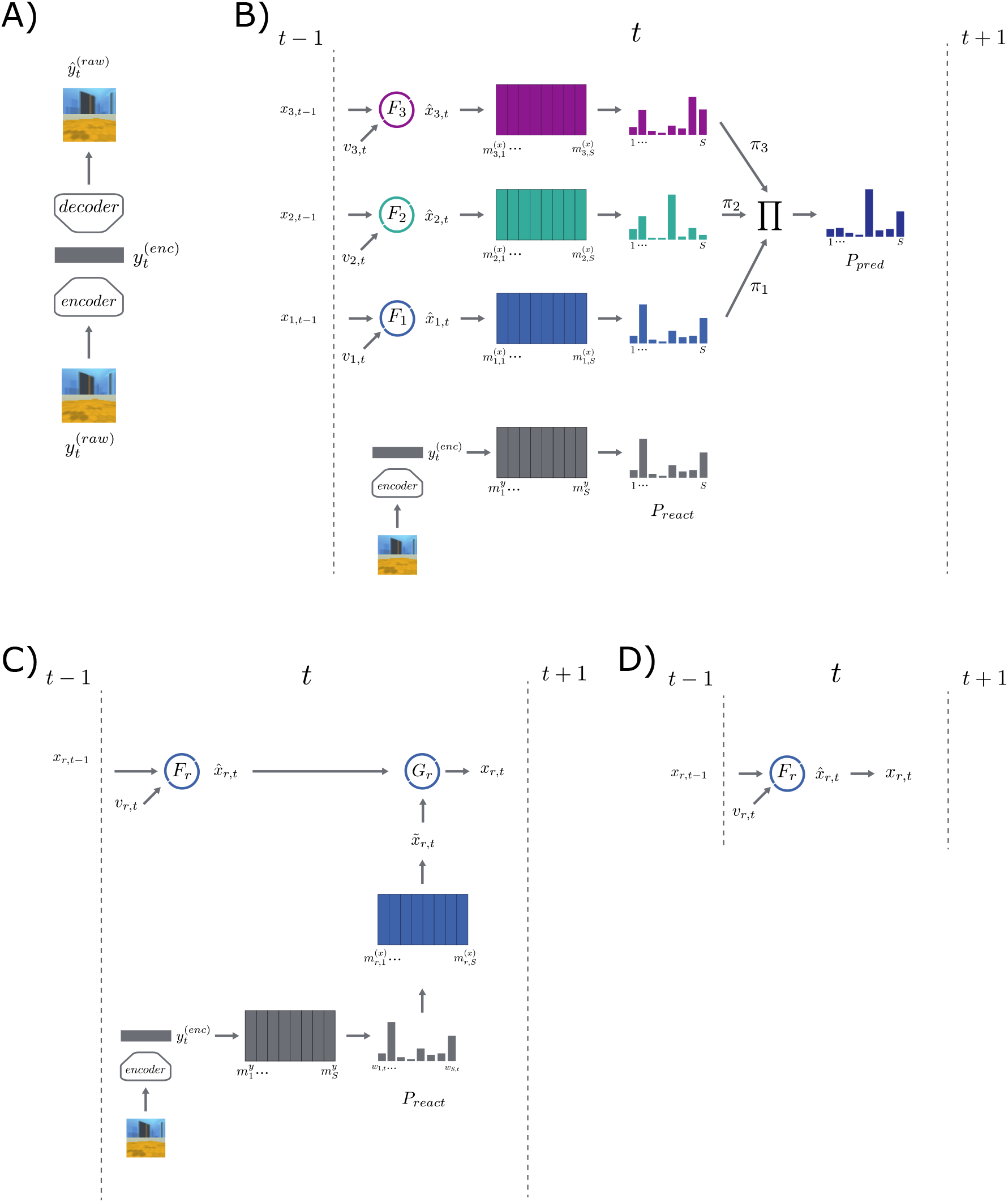
Diagram of the *Spatial Memory Pipeline*. **A)** Computational diagram showing the encoder and decoder networks of the convolutional autencoder used during its training. **B)** Computational diagram showing how the loss is calculated at every time step for an example model with three predictive RNNs. *P*_*react*_ is the target distribution and *P*_*pred*_ the predicted distribution over memories. **C)** Computational diagram of the model dynamics for a time step with visual correction (10% of steps). A correction code 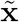 is calculated as a visually-dependent weighted average of visual memory embeddings 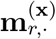. **D)** Computational diagram of the model dynamics for a time step without visual correction (90% of steps).

**Supplementary Figure 2:**
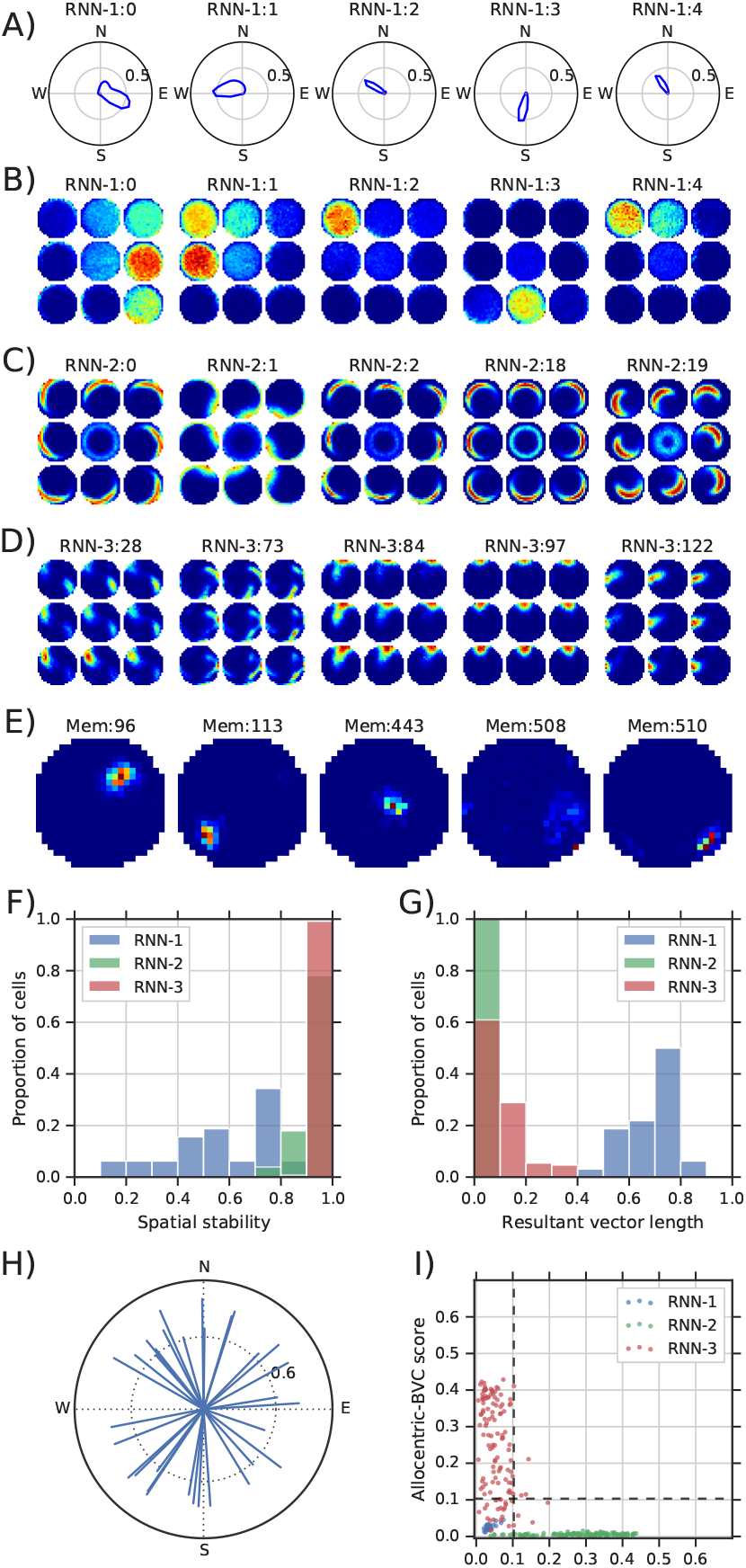
Spa-t12i0al representat12i0ons in the *Spatial Memory Pipeline* trained in a circular open field environment w-1i5t0h wBachk it1e50 walls. **A)** Polar pSlot of activity by heading direction of five cells classified as HD cells. **B)** Spatial ratemaps of five cells shown in A. Central plot shows average activation for each location across all head directions. The eight plots around each central plot show location-specific activity restricted to times when the heading direction of the agent was in the corresponding 45° range (e.g. plot located above the central plot shows average activity when the agent is facing in the north direction). **C)** Spatial ratemaps of five cells classified as egocentric-boundary cells (egoBVCs). **D)** Spatial ratemaps of five cells classified as boundary-vector cells (BVCs). **E)** Spatial ratemaps of five memory slots reactivated by RNN-3. **F)** Spatial stability of cells in each RNN (see Supplementary Methods). **G)** Resultant vector length of cells in each RNN. **H)** Resultant vector of each cell in RNN-1. **I)** Comparison of cells responses to egocentric versus allocentric boundaries in each RNN.

**Supplementary Figure 3:**
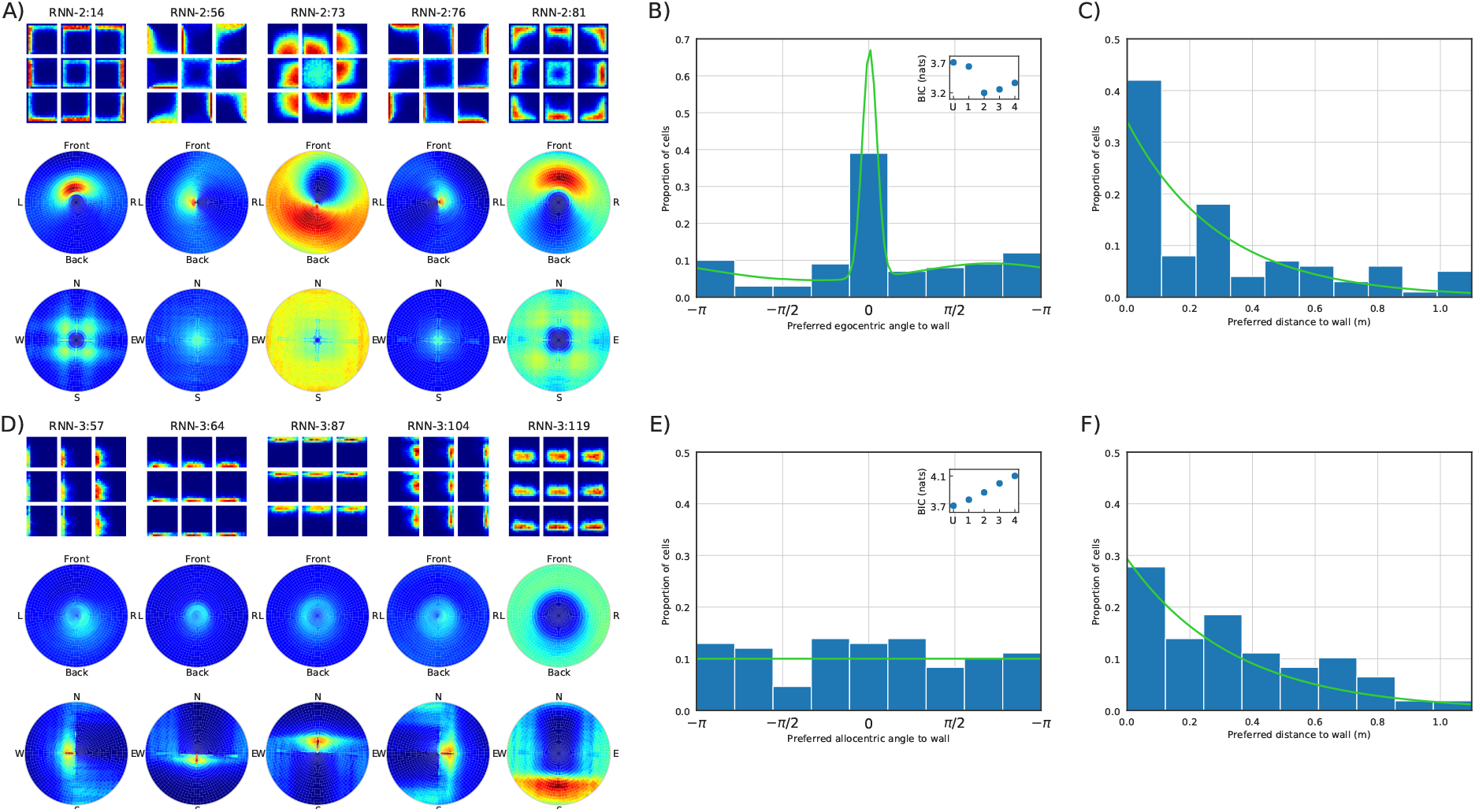
Detail of boundary cells in Fig 2. **A)** First row, ratemap of five cells classified as egoBVCs. Second row, egocentric-boundary ratemap (see Supplementary Methods). Third row, allocentric-boundary ratemap (see Supplementary Methods). **B)** Histogram of preferred egocentric direction to boundary of all units classified as egoBVCs. Inset, Bayes information criterion numbers for a uniform distribution over angles (U) and mixtures of Von Mises distributions with different numbers of components. A mixture of two Von Mises distributions (shown in green), with a mode for boundaries in front agent, was selected (lowest BIC) (see Supplementary Methods). **C)** Histogram of preferred distance to boundary of all units classified as egoBVCs. The distribution over distances was well fit by an exponential distribution (shown in green) with mean distance of 35 cm. **D)** First row, ratemap of five cells classified as (allocentric) BVCs. Second row, egocentric-boundary ratemap. Third row, allocentric-boundary ratemap. **E)** Histogram of preferred allocentric direction to boundary of all units classified as BVCs. Inset shows Bayes information criterion numbers for a uniform distribution over angles (U) and mixtures of Von Mises distributions with different numbers of components. A uniform distribution (shown in green) was selected (lowest BIC). **F)** Histogram of preferred distance to boundary of all units classified as BVCs. The distribution over distances was well fit by an exponential distribution (shown in green) with mean distance of 29 cm.

**Supplementary Figure 4:**
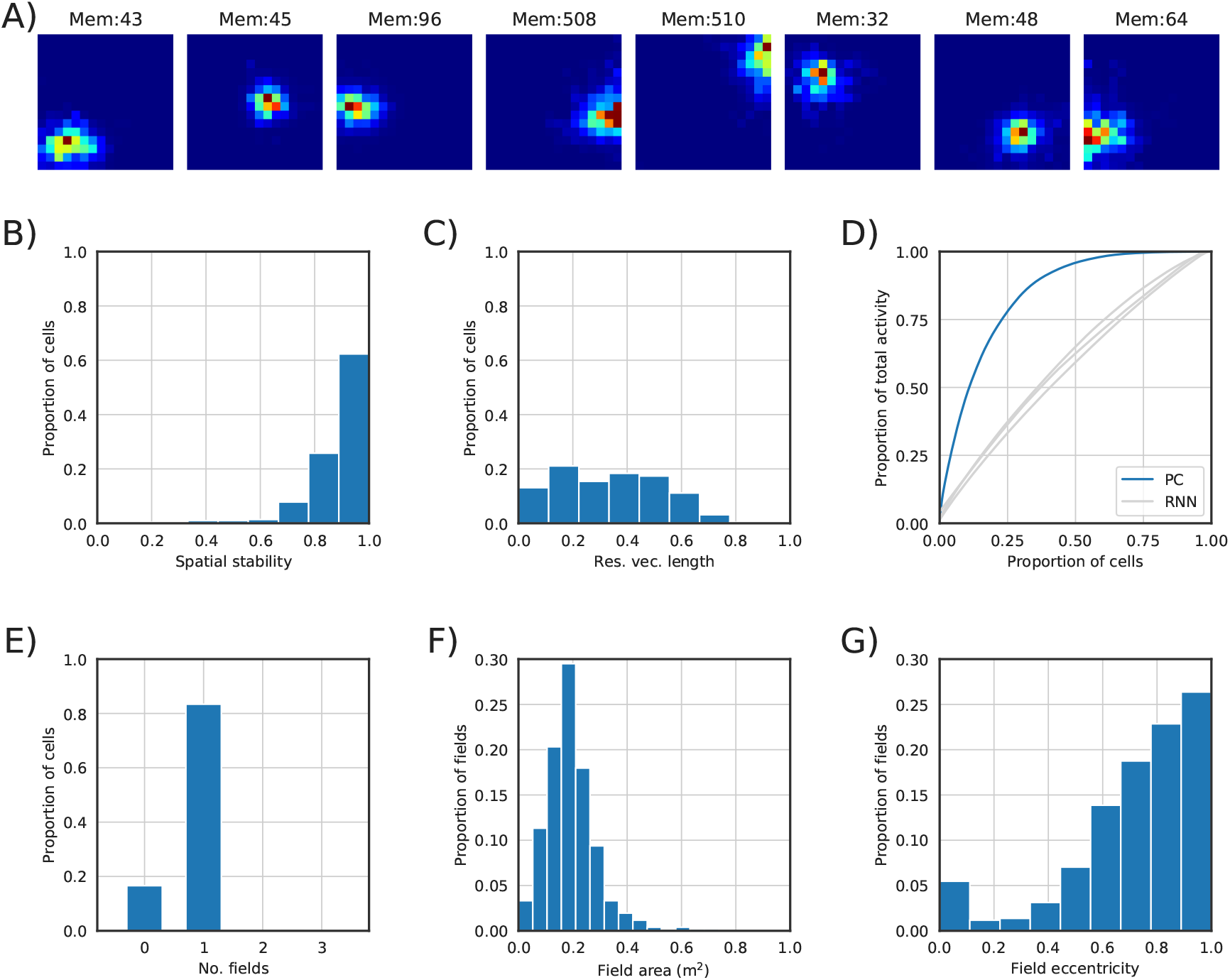
Reactivation of memory slots (containing past RNN-3 states) resembles place-cell responses. **A)** Spatial ratemap of the reactivation of eight memory slots in the experiment described in Fig 2. Activity resembles that of biological place cells in the hippocampus. **B)** Spatial stability of memory slot reactivations (see Supplementary Methods). **C)** Resultant vector of memory slot reactivations. **D)** Cumulative total activity of place cells (memory slot reactivations) ordered by their total activity (blue). Place cells showed a high sparsity: 25% of cells account for 75% of activity. Activity in the three RNNs shows almost no sparsity (cells activated equally on average). **E)** Number of fields per place cell (see Supplementary Methods). **F)** Area of each activity field in square meters (total environment area is 4.8 *m*^2^). **G)** Eccentricity of fields. A score of 1.0 indicates a perfectly round field; lower scores indicate elongation.

**Supplementary Figure 5:**
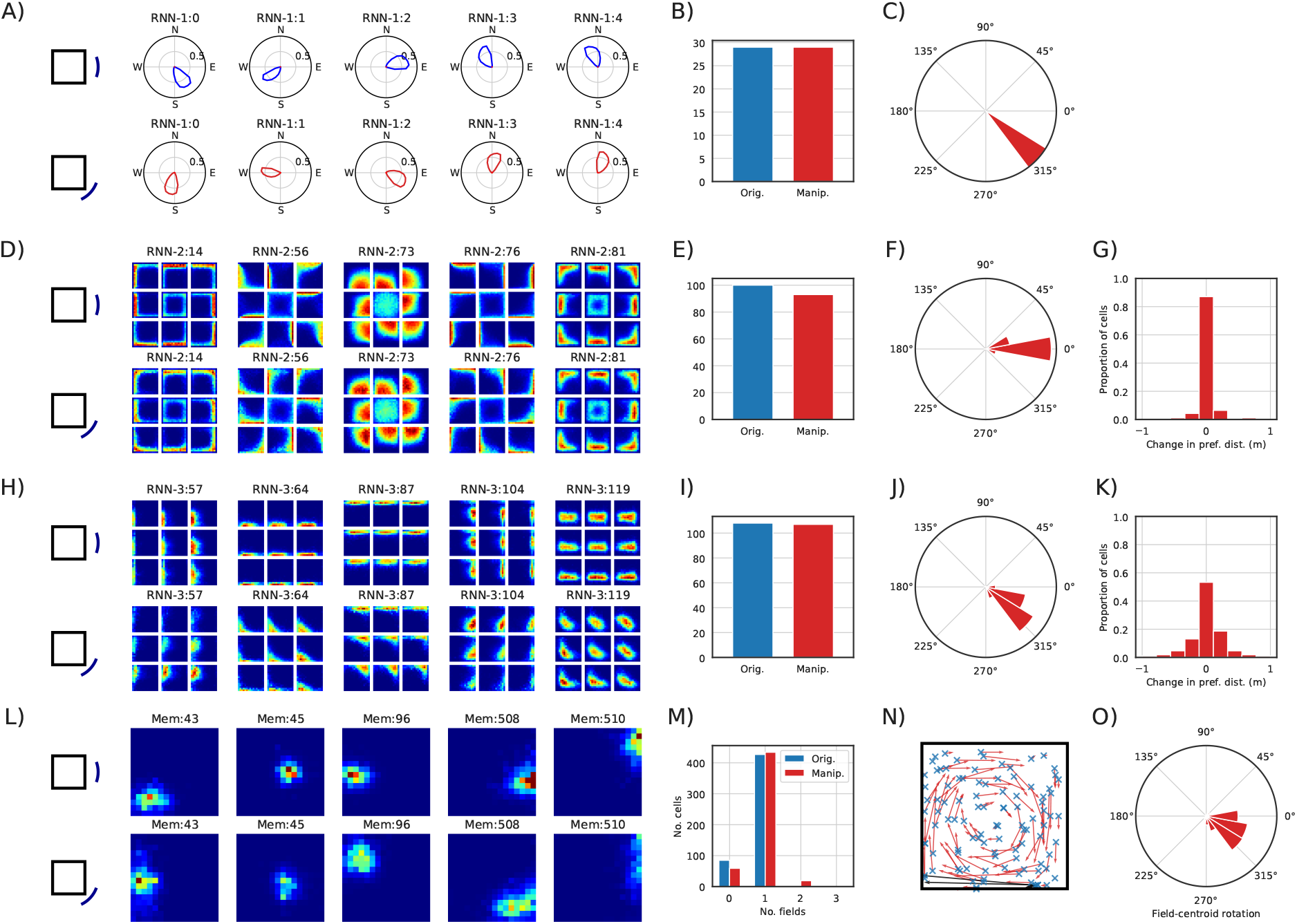
Effects of distal cue rotation (45°) on the representations of the model shown in Fig 2. There was no training after the manipulation. **A)** Polar plots of five cells identified as HD cells. Top row, original environment. Second row, after manipulation. HD activity followed the rotation of distal cues. **B)** Number of RNN units identified as HD cells before and after the environment manipulation. **C)** Histogram of change in phase of the resultant vectors of all HD cells. **D)** Ratemap of five cells identified as egoBVCs. Top row, original environment. Second row, after manipulation. Rotation of distal cues did not affect egoBVCs’ activity. **E)** Number of RNN units identified as egoBVCs before and after the environment manipulation. **F)** Histogram of change in preferred egocentric angle to boundary of egoBVCs. **G)** Histogram of change in preferred distance to boundary of egoBVCs. **H)** Ratemap of five cells identified as (allocentric) BVCs. Top row, original environment. Second row, after manipulation. BVCs followed the rotation of distal cues. **I)** Number of RNN units identified as BVCs before and after the environment manipulation. **J)** Histogram of change in preferred allocentric angle to boundary of BVCs. **K)** Histogram of change in preferred distance to boundary of BVCs. **L)** Ratemap of five place-cell-like units. Top row, original environment. Second row, after manipulation. **M)** Number of fields for each place cell before and after the environment manipulation. **N)** Displacement of place-cell field centroid after manipulation. Blue cross indicates the original centroid, red arrows show their displacement. Black arrows show the displacement when a single field in the original environment split into several fields after manipulation. **O)** Histogram of change in phase of the place-cell centroid of activity when calculated in polar coordinates taking as origin the centre of the enclosure.

**Supplementary Figure 6:**
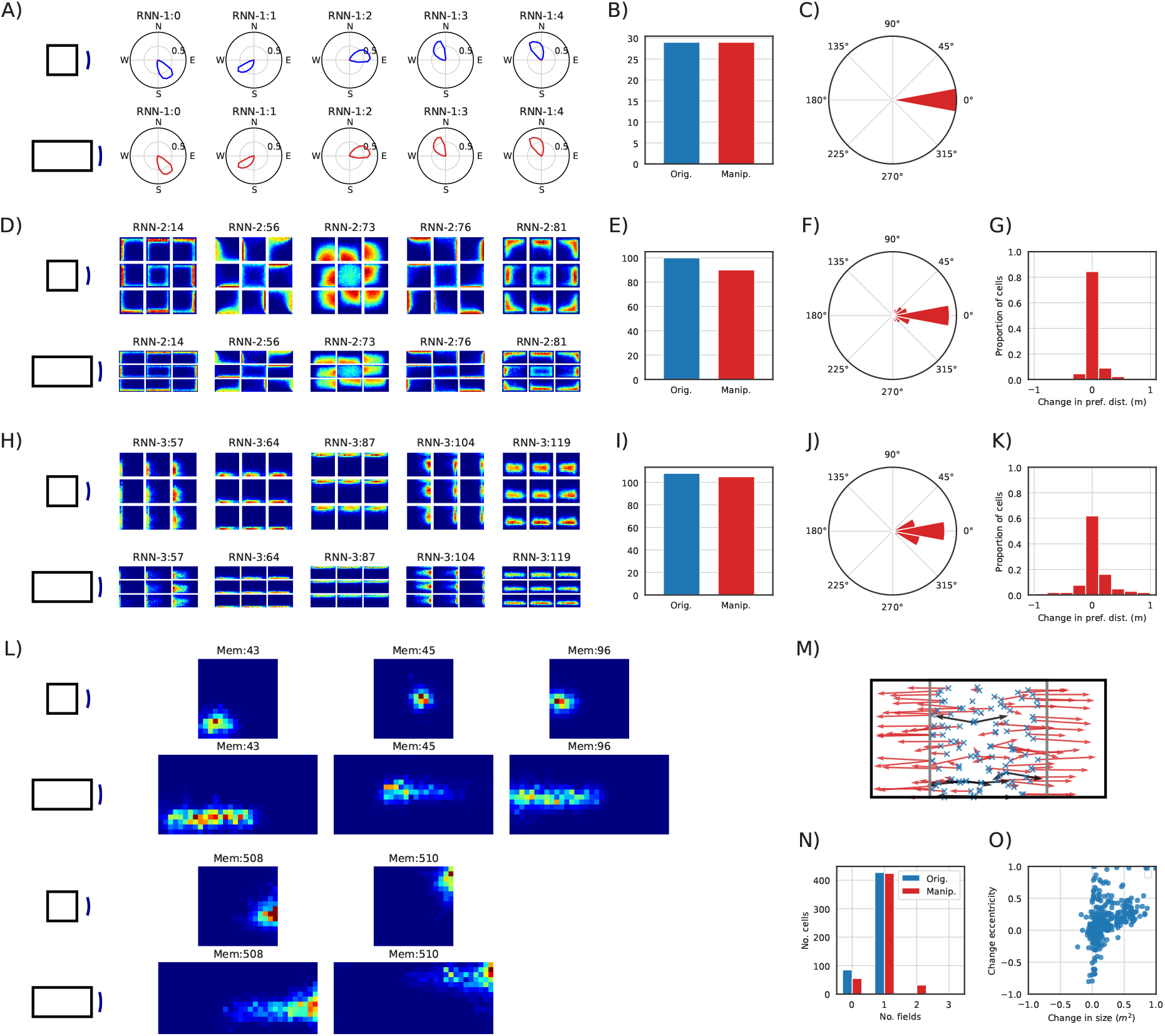
Effects of enclosure resizing (width was doubled) on the representations of the model shown in Fig 2. There was no training after the manipulation. **A)** Polar plots of five cells identified as head-direction cells. Top row, original environment. Second row, after manipulation. The manipulation did not affect HD-cells. **B)** Number of RNN units identified as head-direction cells before and after the environment manipulation. **C)** Histogram of change in phase of the resultant vectors of all HD cells. **D)** Ratemap of five cells identified as egoBVCs. Top row, original environment. Second row, after manipulation. Cells maintained their egocentric response after the stretch. **E)** Number of RNN units identified as egoBVCs before and after the environment manipulation. **F)** Histogram of change in preferred egocentric angle to boundary of egoBVCs. **G)** Histogram of change in preferred distance to boundary of egoBVCs. **H)** Ratemap of five cells identified as (allocentric) BVCs. Top row, original environment. Second row, after manipulation. Cells maintained their allocentric response after the stretch. **I)** Number of RNN units identified as BVCs before and after the environment manipulation. **J)** Histogram of change in preferred allocentric angle to boundary of BVCs. **K)** Histogram of change in preferred distance to boundary of BVCs. **L)** Ratemap of five place-cell-like units. Top, original environment; below, after manipulation. The manipulation caused the activity fields of most cells to stretch with the stretched environment axis. However some cells’ activity field split into two fields. **M)** Displacement of place-cell field centroid after manipulation. Blue cross indicates the original centroid, red arrows show their displacement. Black arrows show the displacement when a single field in the original environment split into several fields after manipulation. **N)** Number of fields for each place cell before and after the environment manipulation. **O)** Change in size and eccentricity of each place cell’s activity field.

**Supplementary Figure 7:**
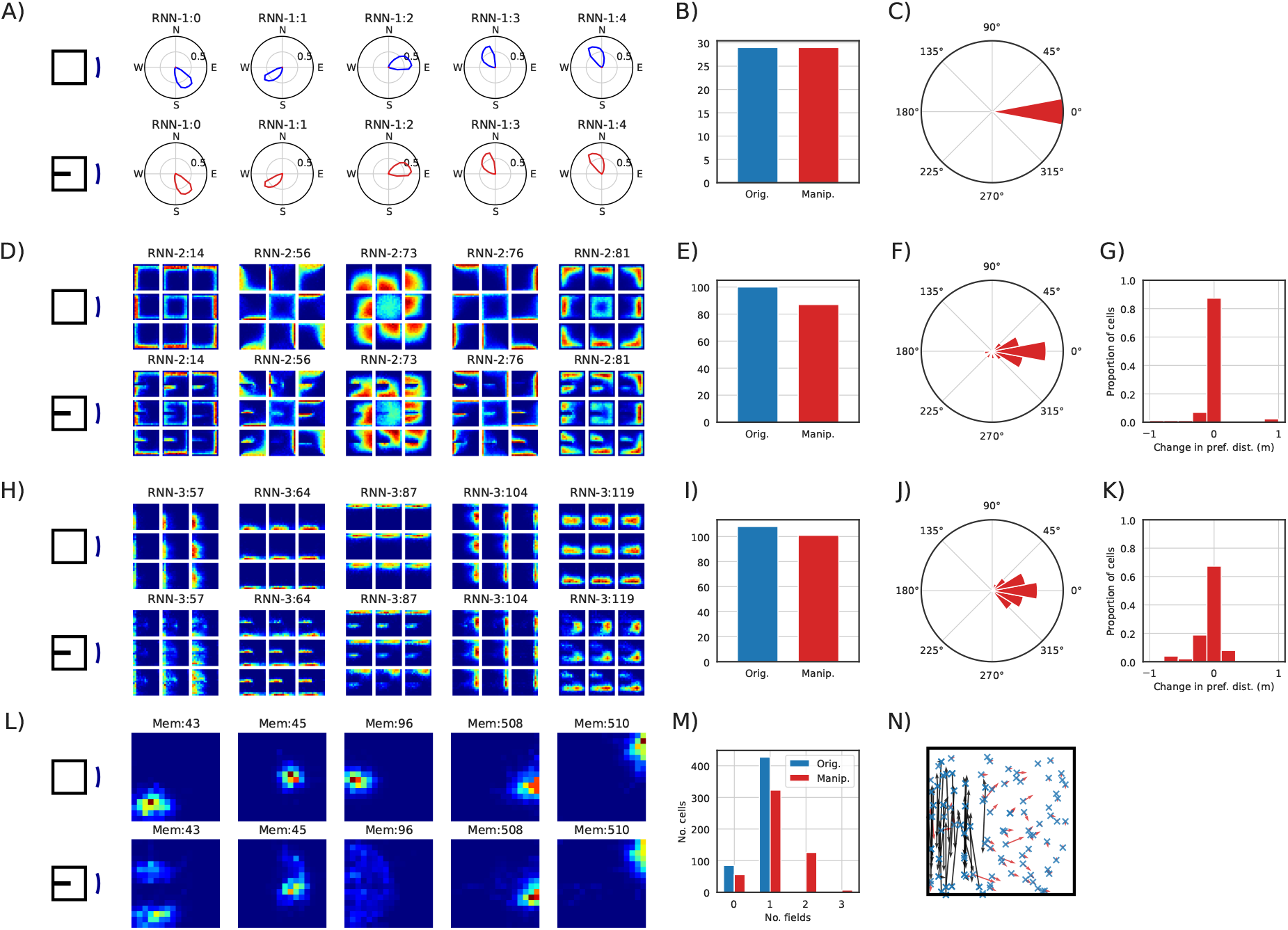
Effects of extra barrier insertion on the representations of the model shown in Fig 2. There was no training after the manipulation. **A)** Polar plots of five cells identified as HD cells. Top row, original environment. Second row, after manipulation. The manipulation did not affect HD cells. **B)** Number of RNN units identified as HD cells before and after the environment manipulation. **C)** Histogram of change in phase of the resultant vectors of all HD cells. **D)** Ratemap of five cells identified as egoBVCs. Top row, original environment. Second row, after manipulation. Cells maintained their egocentric response to the new barrier. **E)** Number of RNN units identified as egoBVCs before and after the environment manipulation. **F)** Histogram of change in preferred egocentric angle to boundary of egoBVCs. **G)** Histogram of change in preferred distance to boundary of egoBVCs. **H)** Ratemap of five cells identified as (allocentric) BVCs. Top row, original environment. Second row, after manipulation. Cells maintained their allocentric response to the new barrier. **I)** Number of RNN units identified as BVCs before and after the environment manipulation. **J)** Histogram of change in preferred allocentric angle to boundary of BVCs. **K)** Histogram of change in preferred distance to boundary of BVCs. **L)** Ratemap of five place-cell-like units. Top row, original environment. Second row, after manipulation. The extra barrier caused cells in the region to duplicate their fields. Cells with distant activity fields were not affected. **M)** Number of fields for each place cell before and after the environment manipulation. **N)** Displacement of place-cell field centroid after manipulation. Blue cross indicates the original centroid, red arrows show their displacement. Black arrows show the displacement when a single field in the original environment split into several fields after manipulation.

**Supplementary Figure 8:**
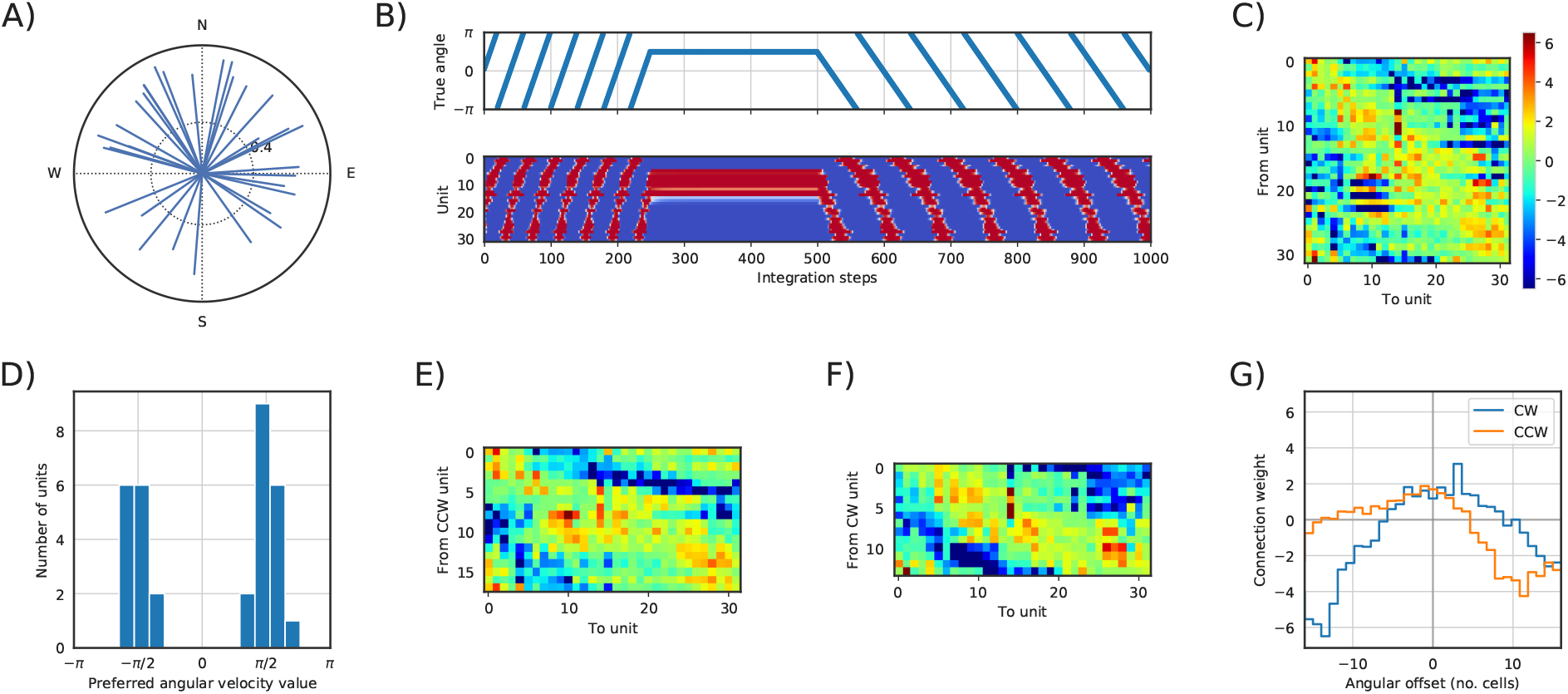
Head direction cell connectivity learnt by a Vanilla-RNN network. **A)** Polar plot of the resultant vectors of cells in RNN-1. **B)** Activation of each unit over 1000 steps of blind integration. Top: True integrated angle. Bottom: Activation of HD-units in RNN-1 through time (units ordered by the phase of their resultant vectors). For the first 250 steps the angular velocity was set to *π*/20 rad/step, the following 250 steps 0 rad/step, the remaining 500 steps *π*/40 rad/step. **C)** Weights in the RNN dynamics matrix, **W**. Columns and rows ordered by the phase of their resultant vector. **D)** Histogram of preferred angular velocities for each cell. **E)** Matrix of weights from CCW cells to all other cells. **F)** Matrix of weights from CW cells to all other cells. **G)** Average weights connecting cells in each of the two rings to all other cells ordered by their angular offset.

**Supplementary Figure 9:**
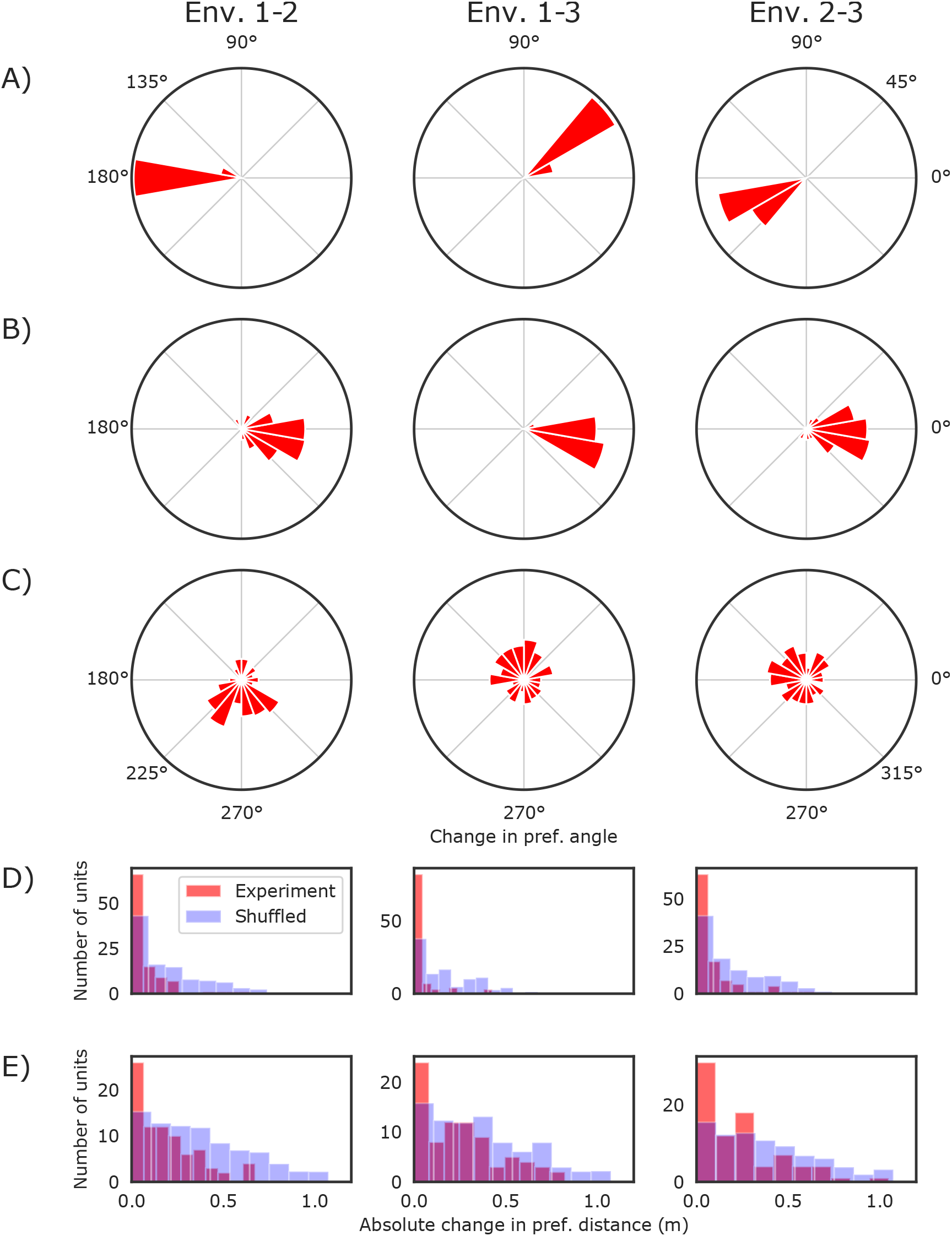
Stability of representations learnt across three environments: square (1), circular (2) and trapezoidal (3). First column compares environments 1 and 2, second column environments 1 and 3, third column environments 2 and 3. **A)** Histogram of HD cell preferred direction differences between environment pairs. HD cells rotate coherently between environments. **B)** Histogram of egoBVC preferred egocentric direction differences between environments. The egoBVCs preserve their angular tuning between environments. **C)** Histogram of BVC preferred allocentric direction differences between environments. The BVCs do not rotate coherently between environments. **D)** Histogram of egoBVC preferred distance differences between environments (red) and normalised histogram of differences between randomly shuffled units (blue). The egoBVCs preserve their distance tuning across environments. **E)** Same as D) for BVCs. The distance tuning is significantly preserved across environments, although not as tightly as in the case of egoBVCs.

**Supplementary Figure 10:**
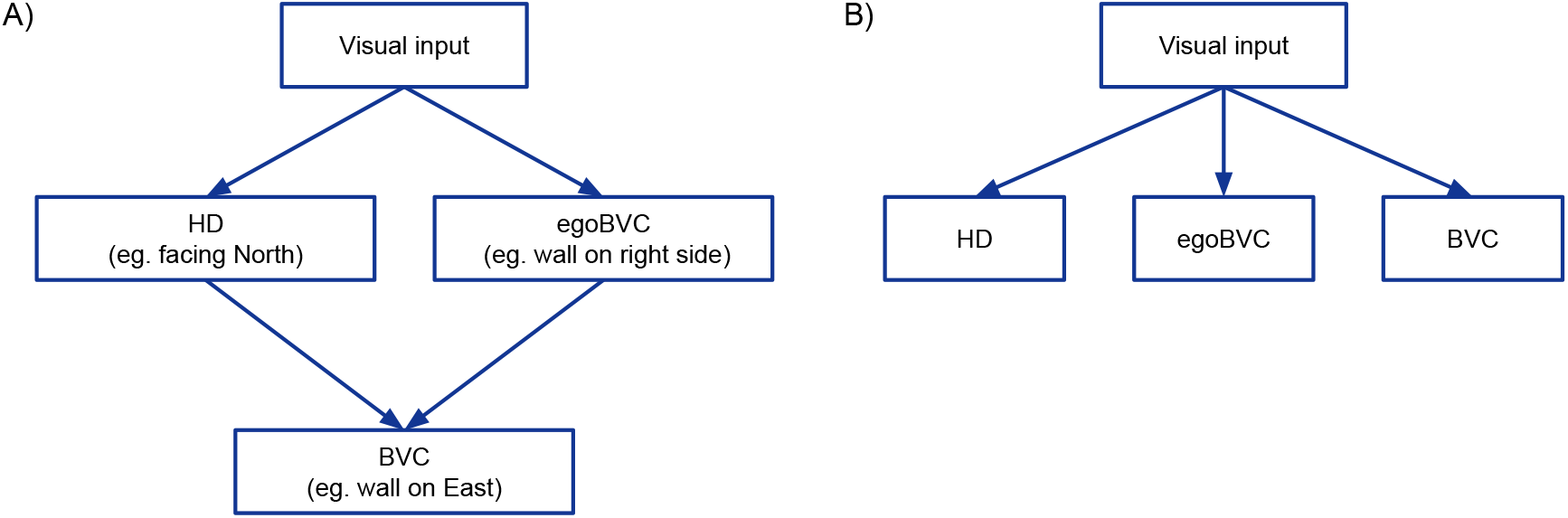
Hypothesised BVC mechanisms. **A)** Traditional BVC mechanism: BVCs result from the conjunction of HD and egoBVC signals. **B)** Proposed BVC mechanism: BVCs are driven by the reactivation of visual memories and temporal coherence.

**Supplementary Figure 11:**
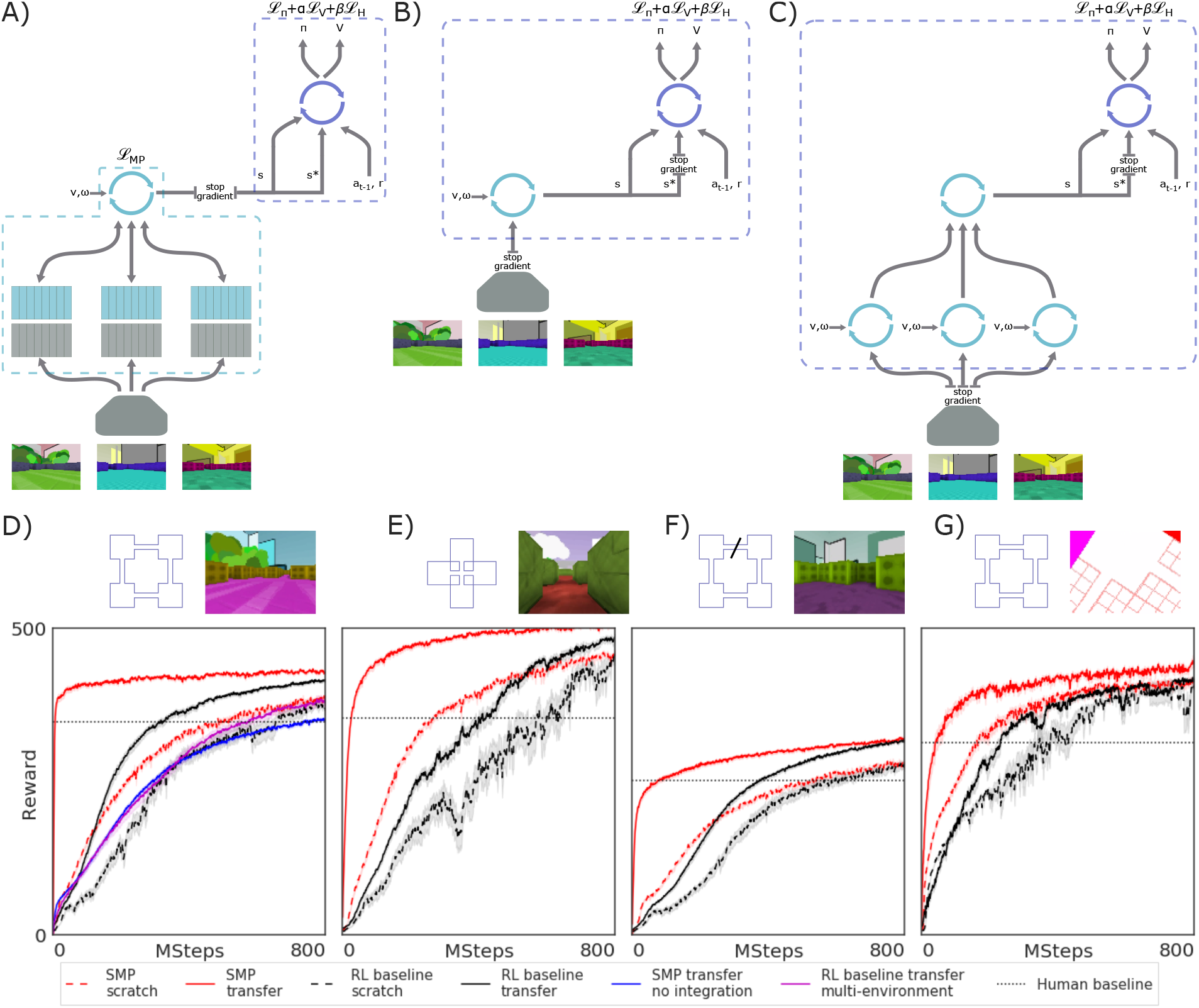
Reinforcement learning architectures and baselines. **A)** Agent training with the *Spatial Memory Pipeline*. The visual inputs from three visually different training environments were stored in separate memory slot banks. The *Spatial Memory Pipeline* (light blue frame) was trained as in the unsupervised experiments to minimise the loss in prediction of memory reactivations. The policy (purple frame) was trained to minimise a combination of policy gradient, value and entropy losses. **B)** Agent training baseline with an LSTM replacing the *Spatial Memory Pipeline*. **C)** Agent training baseline with one LSTM per training environment and a common LSTM replacing the *Spatial Memory Pipeline*. **D-G)** Learning curves in tasks shown in Fig 4C–F. Error regions represent the standard error of the mean episode returns across 5 (D-F) or 1 (G) transfer environment and 25 trained agents (transfer) or 5 seeds (scratch). Red, *Spatial Memory Pipeline*; black, RL baseline. Dashed, learning from scratch; solid, transferred from trained agent. In D), additionally, transfer curves for the training baseline with one LSTM per training environment (magenta), and for a *Spatial Memory Pipeline* with correction at every time step (blue).

**Supplementary Figure 12:**
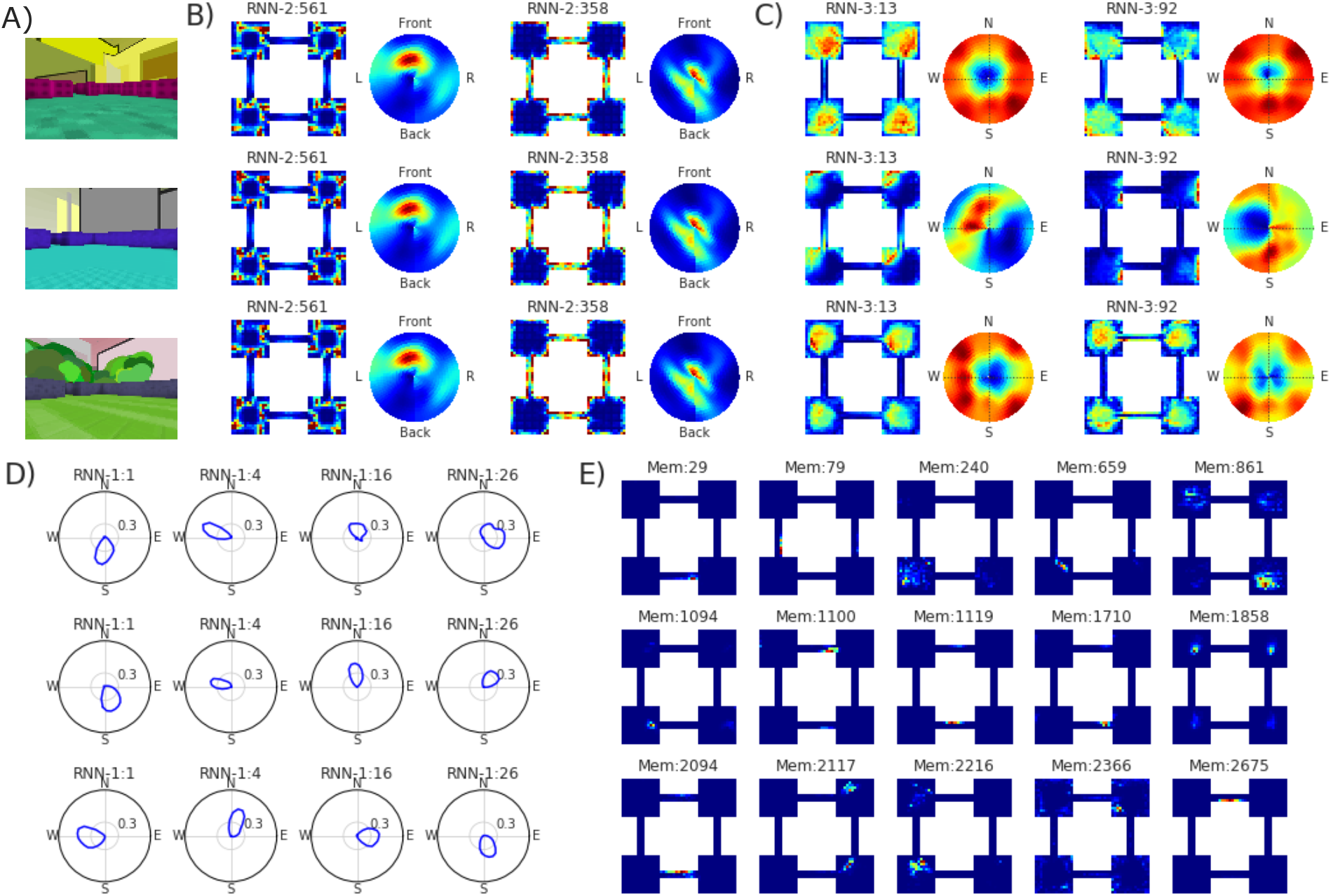
Representations across three reinforcement learning training enclosures, one enclosure in each row. **A)** Agent view of the enclosures. **B)** Ratemaps of two egoBVCs in RNN-2, with their egocentric-boundary ratemap (see Supplementary Methods). **C)** Ratemaps of two BVCs in RNN-3, with their allocentric-boundary ratemap (see Supplementary Methods). **D)** Polar plots of four HD cells in RNN-1. **E)** Ratemaps of five place cells in each environment. Note that, unlike the egoBVC, BVC or HD units, there is no relationship between the place cell units across environments, since they correspond to separate memory banks.

**Supplementary Table 1:** Hyperparameters of rat-like motion model^32^ to generate trajectories for unsupervised experiments.

| Configuration | Value | Meaning |
| --- | --- | --- |
| $b_m$ | 0.13 | Linear velocity Rayleigh scale while moving (m/s) |
| $b_s$ | 0.0 | Linear velocity Rayleigh scale while stopped (m/s) |
| $\mu^{(\phi)}$ | 0 | Rotation velocity Gaussian distribution mean (deg/s) |
| $\sigma^{(\phi)}$ | 330 | Rotation velocity Gaussian distribution standard deviation (deg/s) |
| **d** | 0.03 | Distance to wall to activate wall avoidance (m) |
| **a** | 90 | Angle limit to activate wall avoidance (deg) |
| **slowdown** | 0.25 | Velocity reduction factor when avoidance activated |
| **dt** | 0.02 | Simulation step time increment (s) |
| $\mu_{(stop)}$ | 50 | Mean period for stop/move switching (simulation steps) |
| **n** | 7 | Number of simulation steps per trajectory sample |
| **N** | 500 | Number of samples per trajectory |

**Supplementary Table 2:** Hyperparameters for training in unsupervised experiments.

| Hyperparameter | Single-environment | Multi-environment |
| --- | --- | --- |
| BPTT unroll length | 50 | 50 |
| Batch size | 32 | 32 |
| Training batches | 200,000 | 200,000 |
| Optimiser | **Adam** | **RMSProp** |
| Learning rate (RNNs) | $3 \cdot 10^{-3}$ | $3 \cdot 10^{-4}$ |
| Learning rate (memory embeddings) | $1 \cdot 10^{-2}$ | $3 \cdot 10^{-2}$ |
| Other optimiser params | $\beta_1 = 0.9, \beta_2 = 0.999$ | **decay=0.9, momentum=0** |
| Dropout in RNN states predicting memories | 0.5 | 0.5 |

**Supplementary Table 3:** Hyperparameters for training in reinforcement learning experiments. The learning rate for the actor-critic loss was 1.10^−4^ for all experiments except the top-down schematic representation with RL-baseline, where a slower learning rate of 5.10^−5^ was needed to curb instability.

| Hyperparameter | Value |
| --- | --- |
| BPTT unroll length | 100 |
| Batch size | 64 |
| Optimiser (all losses) | Adam |
| Learning rate (actor-critic loss) | $1 \cdot 10^{-4}$ or $5 \cdot 10^{-5}$ |
| Baseline loss coefficient | 0.5 |
| Entropy loss coefficient | $1 \cdot 10^{-3}$ |
| Learning rate (RNNs) | $1 \cdot 10^{-3}$ |
| Learning rate (memory embeddings) | $3 \cdot 10^{-3}$ |
| Other optimiser params (all losses) | $\beta_1 = 0.9, \beta_2 = 0.999$ |
| Dropout in RNN states predicting memories | 0.5 |

**Supplementary Table 4:**
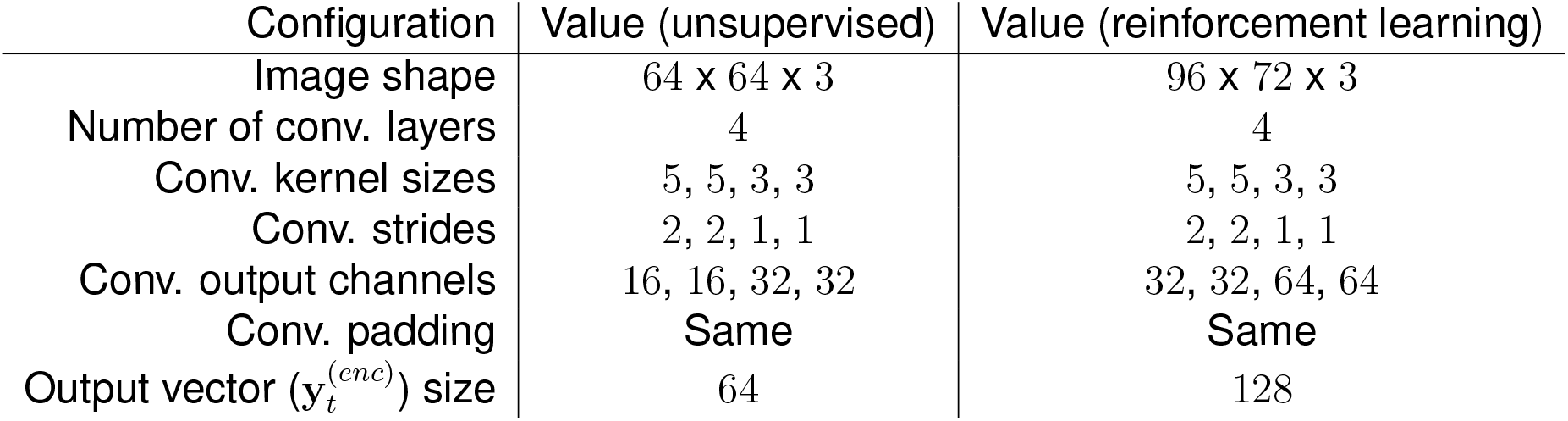
Configuration of the visual encoding level of the *Spatial Memory Pipeline* in unsupervised, and reinforcement learning experiments.

**Supplementary Table 5:** Configuration of the first integration level of the *Spatial Memory Pipeline* in unsupervised and reinforcement learning experiments. Where 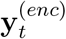 is the ofuttphuet visual encoder, *ω*_*t*_ is the angular velocity, *s*_*t*_ speed parallel to the direction of heading, and (only for reinforcement learning experiments) *s*_⊥,*t*_ the speed perpendicular to the direction of heading.

| Configuration | Value (unsupervised) | Value (reinforcement learning) |
| --- | --- | --- |
| $S$ | 512 | 1024 |
| $R$ | 3 | 3 |
| $\mathbf{y}_t$ | $\mathbf{y}_t^{(enc)}$ | $\mathbf{y}_t^{(enc)}$ |
| $F_1, F_2, F_3$ | Sigmoid-LSTM | Sigmoid-LSTM |
| $G_1, G_2, G_3$ | Sigmoid-LSTM | Sigmoid-LSTM |
| $N_1$ | 32 | 32 |
| $N_2$ | 128 | 768 |
| $N_3$ | 128 | 128 |
| $\mathbf{v}_{1,t}$ | $10 \cdot (\cos(\omega_t), \sin(\omega_t))$ | $10 \cdot (\cos(\omega_t), \sin(\omega_t))$ |
| $\mathbf{v}_{2,t}$ | $10 \cdot (\cos(\omega_t), \sin(\omega_t), s_t)$ | $10 \cdot (\cos(\omega_t), \sin(\omega_t), s_t, s_{\perp,t})$ |
| $\mathbf{v}_{3,t}$ | $()$ | $()$ |
| $H_{react}$ | 0.5 nats | 1.0 nats |
| $P_{correction}$ | 0.1 | 0.1 |
| $P_{storage}$ | 0.0000625 | 0.0001 |

